# Silencing *PHOSPHOENOLPYRUVATE CARBOXYLASE1* in the Obligate Crassulacean Acid Metabolism Species *Kalanchoë laxiflora* causes Reversion to C_3_-like Metabolism and Amplifies Rhythmicity in a Subset of Core Circadian Clock Genes

**DOI:** 10.1101/684050

**Authors:** Susanna F. Boxall, Nirja Kadu, Louisa V. Dever, Jana Kneřová, Jade L. Waller, Peter J. D. Gould, James Hartwell

## Abstract

Unlike C_3_ plants, Crassulacean acid metabolism (CAM) plants fix CO_2_ in the dark using phosphoenolpyruvate carboxylase (PPC; EC 4.1.1.31). PPC combines PEP with CO_2_ (as HCO_3_^−^), forming oxaloacetate that is rapidly converted to malate, leading to vacuolar malic acid accumulation that peaks phased to dawn. In the light period, malate decarboxylation concentrates CO_2_ around RuBisCO for secondary fixation. CAM mutants lacking PPC have not been described. Here, RNAi was employed to silence CAM isogene *PPC1* in *Kalanchoë laxiflora*. Line *rPPC1-B* lacked *PPC1* transcripts, PPC activity, dark period CO_2_ fixation, and nocturnal malate accumulation. Light period stomatal closure was also perturbed, and the plants displayed reduced but detectable dark period stomatal conductance, and arrhythmia of the CAM CO_2_ fixation circadian rhythm under constant light and temperature (LL) free-running conditions. By contrast, the rhythm of delayed fluorescence was enhanced in plants lacking *PPC1*. Furthermore, a subset of gene transcripts within the central circadian oscillator were up-regulated and oscillated robustly. The regulation guard cell genes involved controlling stomatal movements was also altered in *rPPC1-B*. This provided direct evidence that altered regulatory patterns of key guard cell signaling genes are linked with the characteristic inverse pattern of stomatal opening and closing during CAM.

## INTRODUCTION

Crassulacean acid metabolism (CAM) is a pathway of photosynthetic CO_2_ fixation found in species adapted to low rainfall and/ or periodic drought, such as the Madagascan-endemic succulent, *Kalanchoë laxiflora* Baker (Family: Crassulaceae; Order: Saxifragales) (Hartwell et al., 2016). CAM species open their stomata for primary atmospheric CO_2_ fixation in the dark, when their environment is cooler and more humid, and close stomata in the light, when the atmosphere is at its hottest and driest (Osmond, 1978). The increased water use efficiency (WUE) and CO_2_ fixation efficiency of CAM species has led to the proposal that productive CAM crop species, including certain *Agave* and *Opuntia* species, represent a viable approach to generate biomass for biofuels and renewable platform chemicals for industry through their cultivation on under-utilised, seasonally-dry lands that are not well-suited to food crop production (Borland et al., 2009; Cushman et al., 2015). Furthermore, efforts are underway to engineer CAM into key C_3_ crops (Borland et al., 2014; DePaoli et al., 2014; Borland et al., 2015; Lim et al., 2019).

Recently, genome, transcriptome, proteome, and metabolome datasets for a phylogenetically diverse range of independent CAM origins have started to open up a large catalogue of putative CAM genes. CAM species represented by published ‘omics datasets span orchids (*Phalaenopsis equestris* and *Erycina pusilla*, Orchidaceae, monocots), pineapple (*Ananas comosus*, Bromeliaceae, monocots), *Agave* (*A. americana*, *A. deserti*, *A. tequilana*, Agavaceae, monocots), *Yucca* (*Y. aloifolia*, Asparagaceae, monocots), *Kalanchoë* (*K. fedtschenkoi* and *K. laxiflora*, Crassulaceae, dicots), *Mesembryanthemum* (*M. crystallinum*, Aizoaceae, dicots), and *Talinum* (*T. triangulare*, Portulacaceae, dicots) (Cushman et al., 2008; Gross et al., 2013; Cai et al., 2015; Ming et al., 2015; Abraham et al., 2016; Brilhaus et al., 2016; Hartwell et al., 2016; Yang et al., 2017; Heyduk et al., 2018; Heyduk et al., 2019). The optimal exploitation of CAM will be facilitated through decoding the molecular-genetic blueprint for CAM from these genomes and transcriptomes. In turn, to be fully realised, this opportunity requires functional genomics approaches using transgenic and/ or mutant lines of model CAM species to test and define in detail candidate CAM gene function(s) (Dever et al., 2015; Hartwell et al., 2016; Boxall et al., 2017).

During CAM, primary nocturnal CO_2_ assimilation is catalyzed by phospho*enol*pyruvate carboxylase (PPC), generating oxaloacetate that is rapidly converted to malate and stored in the vacuole as malic acid (Borland et al., 2009). At dawn, malic acid is transported out of the vacuole and malate is decarboxylated in the cytosol, with the released CO_2_ re-fixed by RuBisCO behind closed stomata (Borland et al., 2009). Strict temporal control prevents a futile cycle between the enzymes and metabolite transporters driving malate production in the dark, and those driving malate decarboxylation in the light (Hartwell, 2006). PPC regulation is central to this temporal control. PPC is activated allosterically by glucose 6-phosphate (G6P) and inhibited by malate, aspartate and glutamate (O’Leary et al., 2011). The circadian clock optimizes the timing of the CAM carboxylation and decarboxylation pathways to prevent futile cycling (Wilkins, 1992; Hartwell, 2005).

Temporal optimization of PPC involves protein phosphorylation in the dark period, catalysed by a circadian clock-controlled protein kinase, namely phospho*enol*pyruvate carboxylase kinase (PPCK) (Carter et al., 1991; Hartwell et al., 1996; Hartwell et al., 1999; Taybi et al., 2000; Boxall et al., 2005). Phosphorylated PPC is less sensitive to feedback inhibition by malate, which in turn ensures sustained CO_2_ fixation as malic acid accumulates throughout the dark period (Nimmo et al., 1984; Carter et al., 1991; Boxall et al., 2017). In the light, PPC becomes more sensitive to inhibition by malate due to dephosphorylation by a protein phosphatase type 2A (PP2A), which is not known to be subject to circadian control (Carter et al., 1990).

In CAM species such as *Kalanchoë,* the PEP substrate required for nocturnal atmospheric CO2 fixation by PPC is generated through starch breakdown and glycolysis (Borland et al., 2016). In the model C_3_ model species *Arabidopsis thaliana*, nocturnal degradation of leaf starch begins with the phosphorylation of glucan chains by GLUCAN WATER DIKINASE (GWD) and PHOSPHOGLUCAN WATER DIKINASE (PWD) (Ritte et al., 2006). The phosphorylated glucan chains are then further degraded by ALPHA–AMYLASES (AMYs) and BETA-AMYLASES (BAMs) to maltose and glucose. Nocturnal starch hydrolysis by BAMs is the predominant pathway in C_3_ leaves, with chloroplastic BAM3 being the major BAM isozyme driving nocturnal starch degradation in photosynthetic leaf mesophyll cells of Arabidopsis (Fulton et al., 2008). BAM1, BAM2 and BAM5-9 are not required for nocturnal starch degradation in Arabidopsis leaf mesophyll cells (Santelia and Lunn, 2017). Maltose and glucose are exported from chloroplasts by MALTOSE EXCESS 1 (MEX1) and PLASTIDIC GLUCOSE TRANSPORTER (pGlcT), respectively, with MEX1 being the predominant C_3_ route for carbon export (Smith et al., 2005). In the facultative CAM species *M. crystallinum,* chloroplasts possess transporters for triose phosphate (TPT), G6P (GPT), glucose (pGlcT) and maltose (MEX1) (Neuhaus and Schulte, 1996; Kore-eda et al., 2005; Kore-eda et al., 2013). Chloroplasts from C_3_ *M. crystallinum* exported maltose during starch degradation, whereas chloroplasts isolated from CAM-induced leaves exported predominantly G6P, supporting the proposal that starch was broken down via plastidic starch phosphorylase (PHS1) during nocturnal CO_2_ fixation (Neuhaus and Schulte, 1996).

A further defining characteristic of CAM relates to the characteristic nocturnal stomatal opening and CO_2_ uptake, and light period stomatal closure during malate decarboxylation and peak internal CO_2_ supply (Males and Griffiths, 2017). CAM stomatal control is the inverse of the stomatal regulation observed in C_3_ species (Borland et al., 2014). The opening and closing of stomata is driven by the turgor of the guard cell (GC) pair that surround the stomatal pore. High turgor drives opening and reduction in turgor leads to closure. The increase in turgor during opening is driven by the accumulation of K^+^, Cl^−^ and malate^2−^ ions plus sugars in the GCs (Jezek and Blatt, 2017). Closure of stomata is driven by a reversal of GC ion channels and metabolism, with K^+^ and Cl^−^ being transported out, and metabolites being turned over within the GCs. Stomatal aperture responds to changing light, CO_2_, ABA, solutes and water availability (Zhang et al., 2018).

In addition to its role in CAM and C_4_, PPC performs an anapleurotic function by replenishing tricarboxylic acid cycle intermediates utilized for amino acid biosynthesis (Chollet et al., 1996). PPC also functions in the formation of malate as a counter anion for light period opening in C_3_ GCs, and supports nitrogen fixation into amino acids in legume root nodules (Chollet et al., 1996). The major leaf PPCs in Arabidopsis, which are encoded by *PPC1* (AT1G53310) and *PPC2* (AT2G42600), were found to be crucial for leaf carbon and nitrogen metabolism (Shi et al., 2015). The double *ppc1/ppc2* null mutant accumulated starch and sucrose, and had reduced malate, citrate, and ammonium assimilation, and these metabolic changes led to a severe, growth-arrested phenotype (Shi et al., 2015).

Although PPC catalyses primary CO_2_ fixation in CAM and C_4_ plants, the only reported PPC mutants are for the C_4_ dicot *Amaranthus edulis* and the C_4_ monocot *Setaria viridis* (Dever et al., 1995; Alonso-Cantabrana et al., 2018). Loss of the C_4_ *PPC* isogene in *A. edulis* caused a severe and lethal growth phenotype in normal air, with the homozygous mutant plants only managing to reach flowering and set seed when grown at highly elevated CO_2_ (Dever et al., 1995). In *S. viridis*, the C_4_ *PPC* was reduced to very low levels using RNAi in transgenic lines (Alonso-Cantabrana et al., 2018). These lines grew very slowly even at 2 % CO_2_ (normal air is 0.04 %), and developed increased numbers of plasmodesmatal pit fields at the mesophyll-bundle sheath interface (Alonso-Cantabrana et al., 2018). By contrast, no CAM mutants lacking PPC have been described, but transgenic lines of *Kalanchoë* lacking the light-period, decarboxylation pathway enzymes mitochondrial NAD-malic enzyme (NAD-ME) and pyruvate orthophosphate dikinase (PPDK) displayed a near complete loss of dark CO_2_ fixation and failed to turnover significant malate in the light period (Dever et al., 2015). In addition, a CAM mutant of the inducible CAM species *Mesembryanthemum crystallinum* has been described lacking the starch synthesis enzyme plastidic phosphoglucomutase (Cushman et al., 2008).

In addition, the CAM-associated *PPCK1* gene in *Kalanchoë* was also silenced using RNAi, and this led not only to a reduction in dark period CO_2_ fixation, but also perturbed the operation of the central circadian clock (Boxall et al., 2017). However, even the strongest *PPCK1* RNAi line was still able to acheive ∼33 % of the dark period CO_2_ fixation observed in the wild type (Boxall et al., 2017). Those findings led us here to develop transgenic lines of *K. laxiflora* in which the CAM-associated isogene of *PPC* itself (isogene *PPC1*) was down-regulated using RNAi. The most strongly silenced line, *rPPC1-B,* lacked *PPC1* transcripts and activity, and this resulted in the complete loss of dark CO_2_ fixation associated with CAM, and arrhythmia of the CAM CO_2_ fixation rhythm under constant light and temperature (LL) free-running conditions. Growth of *rPPC1-B* plants was reduced relative to wild type in both well-watered and drought-stressed conditions. The plants reverted to fixing CO_2_ in the light, especially in their youngest leaf pairs. Although the circadian rhythm of CO_2_ fixation dampened rapidly towards arrhythmia in the *rPPC1-B* line, the distinct circadian clock output of delayed fluorescence, and the oscillations of the transcript abundances of a subset of core circadian clock genes, were enhanced. Furthermore, the temporal phasing of a wide range of GC specific signaling genes involved in opening and closing was perturbed relative to the wild type in *rPPC1-B*.

## RESULTS

### Initial Screening and Characterisation of *PPC1* RNAi Lines of *K. laxiflora*

As *K. laxiflora* is relatively slow to grow, flower and set seed, taking around 9-months seed-to-seed (Hartwell et al., 2016), data are presented for independent primary transformants that were propagated clonally via leaf margin plantlets and/ or stem cuttings. Primary transformants were screened initially using high-throughput leaf disc tests for starch and acidity at dawn and dusk (Cushman et al., 2008; Dever et al., 2015). Independent transgenic lines that acidified less during the dark period were screened for the steady-state abundance of *PPC1* transcripts using RT-qPCR. Line *rPPC1-B* displayed a complete loss of *PPC1* transcripts, whereas *rPPC1-A* had an intermediate level of *PPC1* transcripts (Figure 1A). The other plant-type *PPC* genes (*PPC2, PPC3* and *PPC4*) were up-regulated relative to the wild type, with their peak phased to dawn (Figure 1B to 1D). *PPC2* was up-regulated in line *rPPC1-B* at 6 and 10 h into the 12 h dark period (Figure 1B).

**Figure 1.**
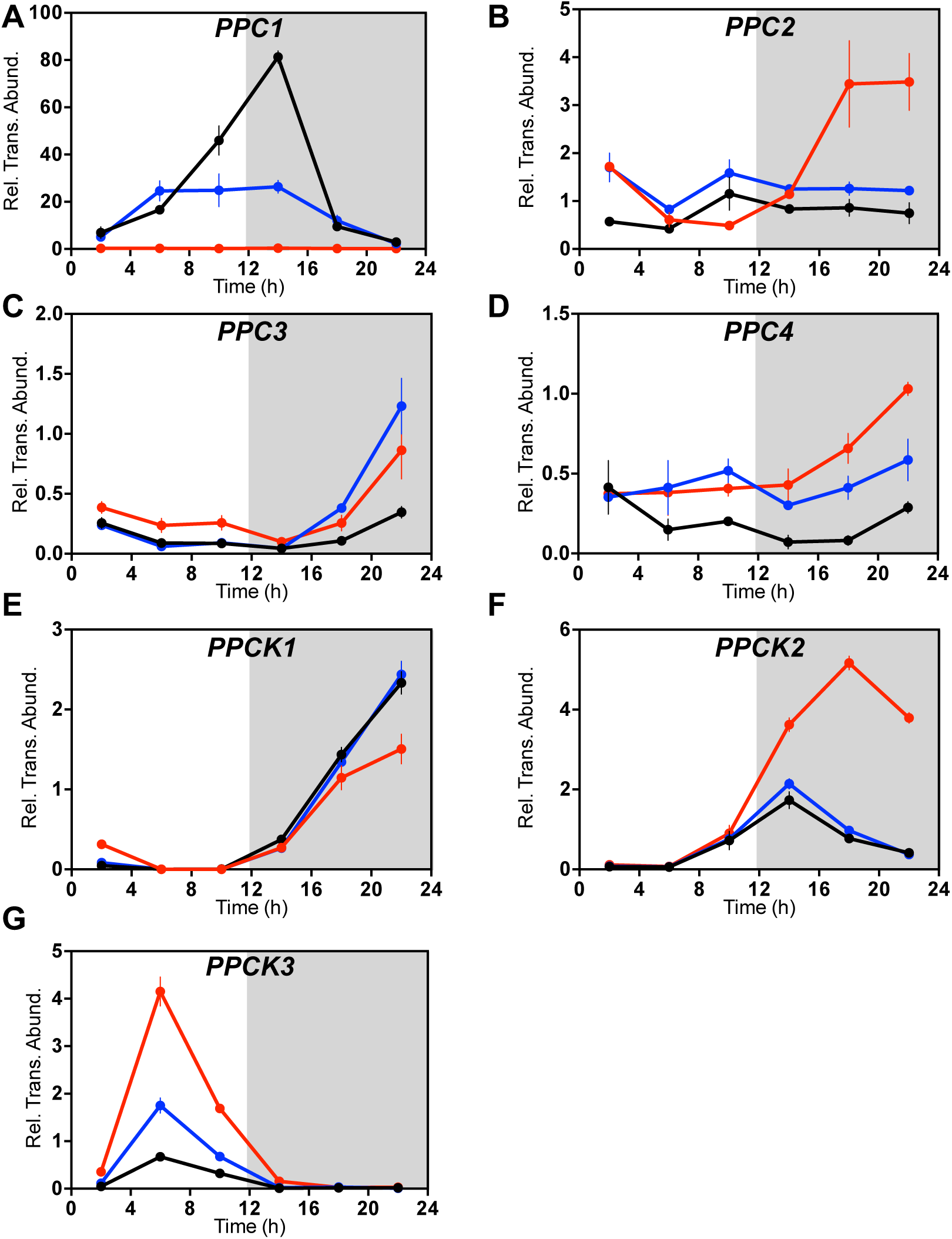
Confirmation of Target Gene Silencing in Transgenic *K. laxiflora* RNAi Lines *rPPC1-A* and *rPPC1-B*. Gene transcript abundance in leaf pair 6 (LP6) was measured using RT-qPCR for target genes: **(A)** *PPC1*; **(B)** *PPC2*; **(C)** *PPC3*; **(D)** *PPC4;* **(E)** *PPCK1;* **(F)** *PPCK2; and* **(G)** *PPCK3*. LP6 were sampled every 4 h across the 12-h-light/ 12-h-dark cycle. A *THIOESTERASE/THIOL ESTER DEHYDRASE-ISOMERASE* superfamily gene (*TEDI*) was amplified from the same cDNAs as a reference gene. Gene transcript abundance data represents the mean of 3 technical replicates for biological triplicates and was normalized to loading control gene *(TEDI)*; error bars represent the standard error. In all cases, plants were entrained under 12-h-light/ 12-h-dark cycles for 7-days prior to sampling. Black data are for the wild type, blue data *rPPC1-A,* and red data *rPPC1-B*.

In line *rPPC1-B*, the transcript level of the protein kinase that phosphorylates the CAM-associated PPC1, namely *PPCK1*, was down-regulated during its dark period phased peak, and slightly up-regulated at 2 h after dawn (Figure 1E). The other two detectable *PPCK* genes, namely *PPCK2* and *PPCK3,* were up-regulated, with *PPCK2* induced 5-fold in *rPPC1-B* when it reached its 24 h peak in the middle of the dark period, 4 h after the wild type peak (Figure 1F). *PPCK3* peaked in the middle of the light, when it reached a level almost 8-fold greater than the wild type (Figure 1G).

### Loss of *PPC1* Transcripts led to Loss of PPC Protein and Activity

Immunoblotting using an antibody raised against purified CAM-specific PPC protein from *K. fedtschenkoi* leaves (Nimmo et al., 1986) revealed reduced PPC protein in *rPPC1-A,* and no detectable PPC in *rPPC1-B* (Figure 2A).

**Figure 2.**
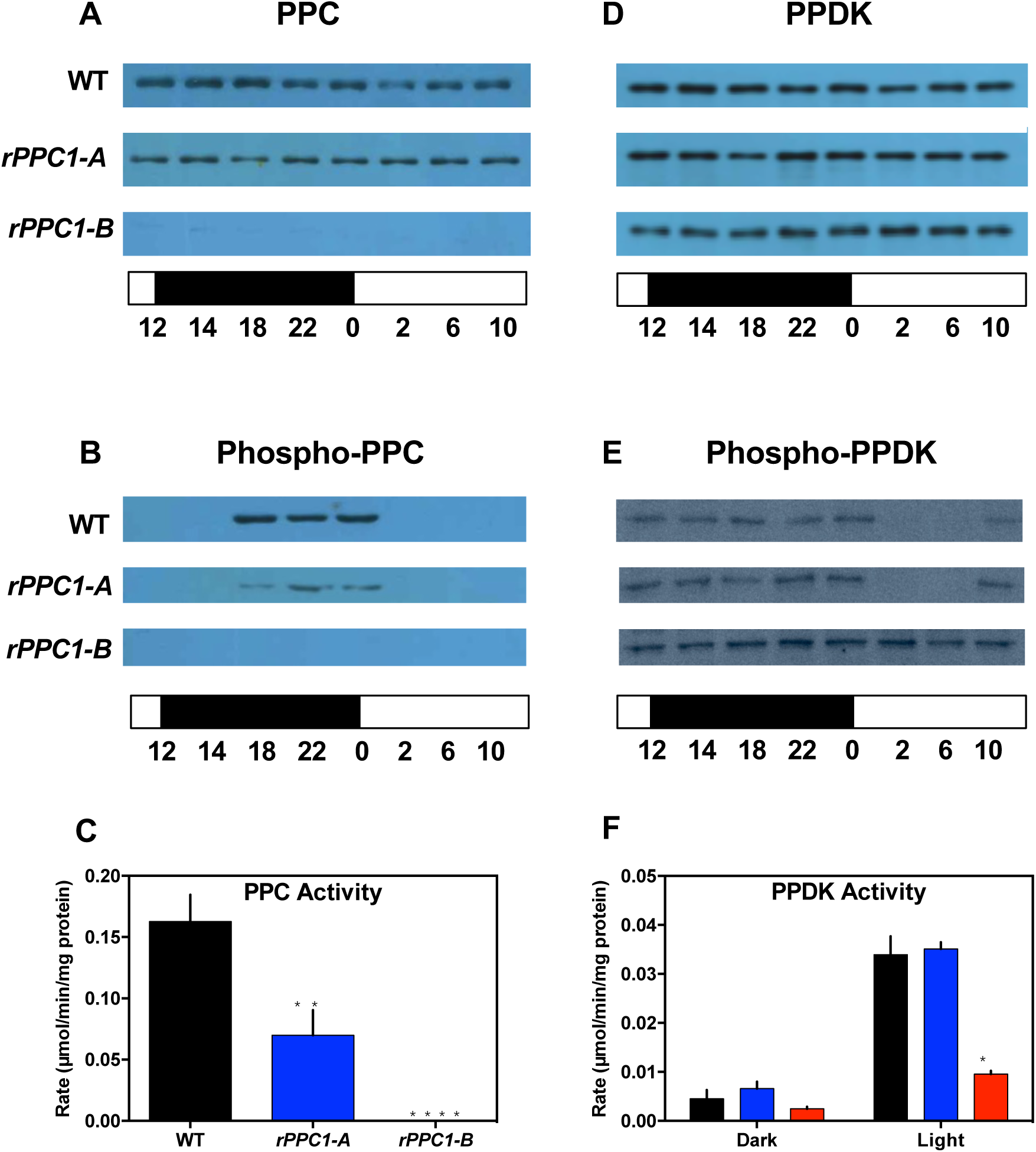
Loss of PPC1 Led to Loss of PPC Protein and Activity and Failure to Activate PPDK in the Light. Total leaf protein from leaf pair 6 (LP6) was isolated from leaves sampled at dawn and dusk plus every 4 h, starting at 2 h into the light, across the 12-h-light/ 12-h-dark cycle, separated using SDS-PAGE and used for immunoblot analyses with antibodies raised to PPC **(A)**, and phospho-PPC **(B)**. **(C)** PPC activity was measured in rapidly desalted extracts from LP6, ** = *rPPC1-A* p= 0.0025*, **** = rPPC1-B,* p<0.0001. **(D)** Anti-PPDK immunoblot. **(E)** Anti-phospho-PPDK immunoblot. **(F)** PPDK activity was measured in leaf extracts prepared from both dark leaves and illuminated leaves*, * = rPPC1-B,* p=0.019. The activities of PPC, measured 6 h into the light period, and PPDK, measured both at 6h into the dark, and 6h into the light, were the mean of 3 technical replicates of 3 biological replicates. The white bar below each panel represents the 12-h-light period and the black bar below each panel represents the 12-h-dark period. Black data are for the wild type, blue data *rPPC1-A,* and red data *rPPC1-B*.

The level of phospho-PPC was also measured using immunoblotting (Figure 2B). Phospho-PPC in *rPPC1-A* was lower than the wild type, whilst *rPPC1-B* lacked detectable phospho-PPC (Figure 2B); consistent with the level of PPC (Figure 2A). Although *PPC2* was up-regulated in *rPPC1-B* (Figure 1B), it was not detected at the protein level by the PPC antibody (Figure 2A). Furthermore, despite *PPCK2* and *PPCK3* being up-regulated by 5- to 8-fold in *rPPC1-B* (Figure 1F and 1G), any protein produced from these transcripts did not phosphorylate any PPC2 protein that may have resulted from the induced *PPC2* transcripts (Figure 2B); at least not within the limits of detection with this immunoblotting technique. Rapidly desalted leaf extracts were used to measure the apparent K_i_ of PPC for L-malate. The K_i_ was higher in the dark than the light for wild type and *rPPC1-A*, but no change in the K_i_ was detected for *rPPC1-B* (Supplemental Figure 1). Furthermore, PPC activity was not detected in rapidly desalted extracts from *rPPC1-B* leaves, whereas *rPPC1-A* displayed reduced but detectable PPC activity, at 43 % of the wild type level (Figure 2C).

Pyruvate orthophosphate dikinase (PPDK), which functions in concert with NAD-ME during the light-period as part of the CAM malate decarboxylation pathway (Dever et al., 2015), was measured on immunoblots using specific antibodies against PPDK and phospho-PPDK (Chastain et al., 2000; Chastain et al., 2002). The blots showed a similar amount of PPDK protein over the diel cycle in wild type, *rPPC1-A* and *rPPC1-B* (Figure 2D). In *Kalanchoë*, PPDK is inactivated in the dark by phosphorylation by PPDK-regulatory protein (PPDK-RP), which also activates PPDK in the light through dephosphorylation (Dever et al., 2015). In the wild type, immunoblotting of phospho-PPDK revealed that PPDK was dephosphorylated, and therefore likely to be fully active, between 02:00 and 06:00 in the light when it is required for conversion of pyruvate, from malate decarboxylation, to PEP, thereby facilitating recycling of carbon through gluconeogenesis to starch (Figure 2E). Line *rPPC1-A* showed the same pattern of PPDK phosphorylation/ de-phosphorylation as the wild type (Figure 2E). However, PPDK was phosphorylated throughout the 24 h cycle in *rPPC1-B* (Figure 2E), and, therefore, likely to be inactive. Consistent with this prediction, loss of the light period dephosphorylation of PPDK in *rPPC1-B* correlated with a significant decrease in PPDK activity in the light, whereas the wild type and line *rPPC1-A* showed strongly light-induced levels of PPDK activity that correlated with the detected level of PPDK dephosphorylation in the light (Figure 2F).

### Levels of Malate, Starch and Soluble Sugars

During CAM in *Kalanchoë*, primary nocturnal fixation of atmospheric CO_2_ (as HCO_3_^−^) results in vacuolar malic acid accumulation throughout the dark period. Starch accumulates in the light period and is broken down during the dark to provide PEP as the substrate for carboxylation by PPC. Starch is also broken down in a rapid burst at dawn to form soluble sugars (Wild et al., 2010; Boxall et al., 2017). As the lack of CAM-associated *PPC1* was predicted to prevent primary nocturnal carboxylation, metabolites including starch, malate and soluble sugars were measured every 4 h over the 24 h cycle (Figure 3).

**Figure 3.**
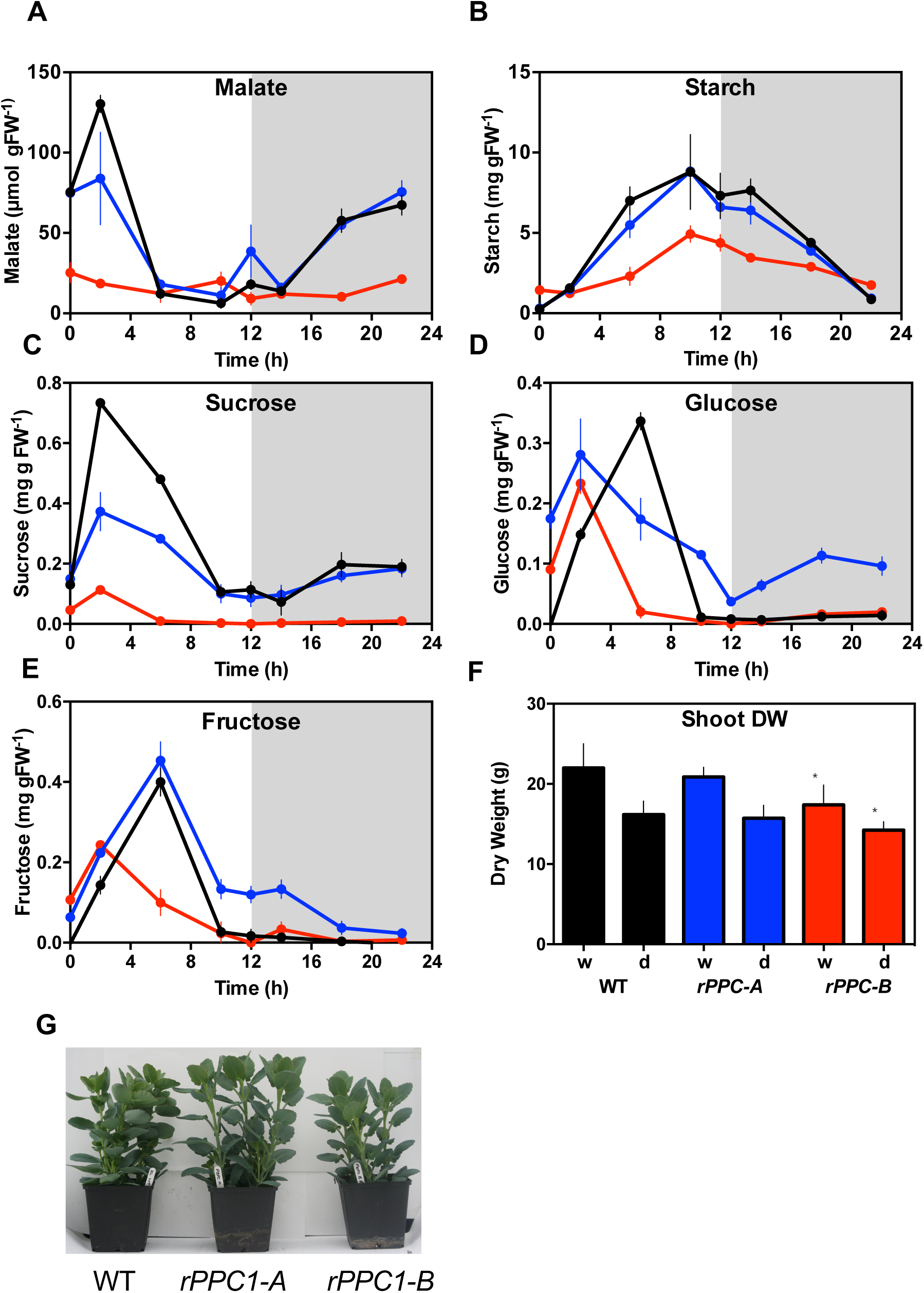
Impact of silencing *PPC* on malate, starch, soluble sugars and growth. **(A)** Malate content, **(B)** starch, **(C)** sucrose, **(D)** glucose and **(E)** fructose were determined from leaf pair 6 (LP6) samples collected every 4 h using plants entrained under 12-h-light/ 12-h-dark cycles. Methanol extracts were prepared from the leaves from wild type, *rPPC1-A* and *rPPC1-B* and used for malate and soluble sugar determination, whereas starch was measured in the insoluble pellet. **(F)** Dry weight of 168 day old greenhouse grown plants, either well-watered (w) or drought-stressed for the last 30 days (d). * denotes significant difference between wild type and *rPPC1-B*. Well-watered *rPPC1-B* p=0.026, drought-stressed *rPPC1-B* p= 0.0063. n = 10 developmentally synchronised, clonal plants per line. **(G)**, 4-month-old plants raised in a glasshouse under 16-h-light/ 8-h-dark. Black data are for the wild type, blue data *rPPC1-A,* and red data *rPPC1-B*.

Wild type plants accumulated 130 µmols gFW^−1^ malate by dawn, whereas *rPPC1-A* and *rPPC1-B* accumulated 75 µmol gFW^−1^ and 19.5 µmols gFW^−1^, respectively (Figure 3A). The Δ-malate values for wild type, *rPPC1-A* and *rPPC1-B* were 124.0, 64.3 and 16.2 µmol gFW^−1^ (Figure 3A). During the diel cycle, *rPPC1-A* and *rPPC1-B* synthesized 100 % and 41 % of the amount of starch accumulated by the wild type (Figure 3B). The Δ-starch values for wild type, *rPPC1-A* and *rPPC1-B* were 8.5, 8.5 and 3.5 mg starch gFW^−1^ (Figure 3B). Lines *rPPC1-A* and *rPPC1-B* accumulated 51 % and 15 % of the amount of sucrose accumulated by the wild type (Figure 3C), and 83 % and 69 % of the level of glucose (Figure 3D). Glucose accumulated 4 h after the sucrose peak in the wild type, whereas glucose levels peaked at the same time as sucrose in lines *rPPC1-A* and *rPPC1-B* (Figure 3C and CD). Finally, lines *rPPC1-A* and *rPPC1-B* accumulated, respectively, 113 % and 61 % of the amount of fructose compared with the wild type (Figure 3E). In *rPPC1-B,* the daily maxima for fructose and glucose occurred coincident with that of sucrose at 2 h after dawn (Figure 3C to 3E).

### Growth Analysis in Well-Watered versus Drought-Stressed Conditions

CAM is widely regarded as an adaptation to drought, and so it was important to compare the growth performance of wild type (full CAM) with that of *rPPC1-A* (small reduction in CAM) and *rPPC1-B* (no CAM) under both well-watered (WW) and drought-stressed (D-S) conditions (Figure 3F and Supplemental Figure S2). *rPPC1-B* were significantly smaller than wild type in both WW and D-S conditions (Figure 3F). In WW conditions, the shoot dry weight of *rPPC1-B* was 21 % less than the wild type (Figure 3F and S2), but *rPPC1-A* was not significantly different to the wild type. Under D-S, the shoot dry weight of line *rPPC1-B* was 12 % lower than that of the wild type (Figure 3F). The visible size of representative 4-month-old plants demonstrated that *rPPC1-B* was smaller than wild type and *rPPC1-A* (Figure 3G), consistent with the shoot dry weight data (Figure 3F). Shoot fresh weight was also lower in WW *rPPC1-B* (Supplemental Figure S2). Furthermore, *rPPC1-B* displayed a reduced degree of leaf in-rolling in response to drought relative to the wild type (Supplemental Figure S3).

### Gas Exchange Characteristics under Light/ Dark Cycles

The gas exchange of mature CAM leaves (leaf pair 6, LP6) of each line was measured over a 12-h-light, 25°C, 60 % humidity/ 12-h-dark, 15°C, 70 % humidity cycle using an infra-red gas analyser (IRGA; LI-COR LI-6400XT) (Figure 4A to 4C). Over the 24 h diel cycle, wild type fixed 297 µmol CO_2_ m^−2^, *rPPC1-A* 320 µmol CO_2_ m^−2^, and *rPPC1-B* 230 µmol CO_2_ m^−2^ (Figure 4A). Net CO_2_ fixation was 23 % less than the wild type for *rPPC1-B*, whereas line *rPPC1-A* was 15 % greater (Figure 4A).

**Figure 4.**
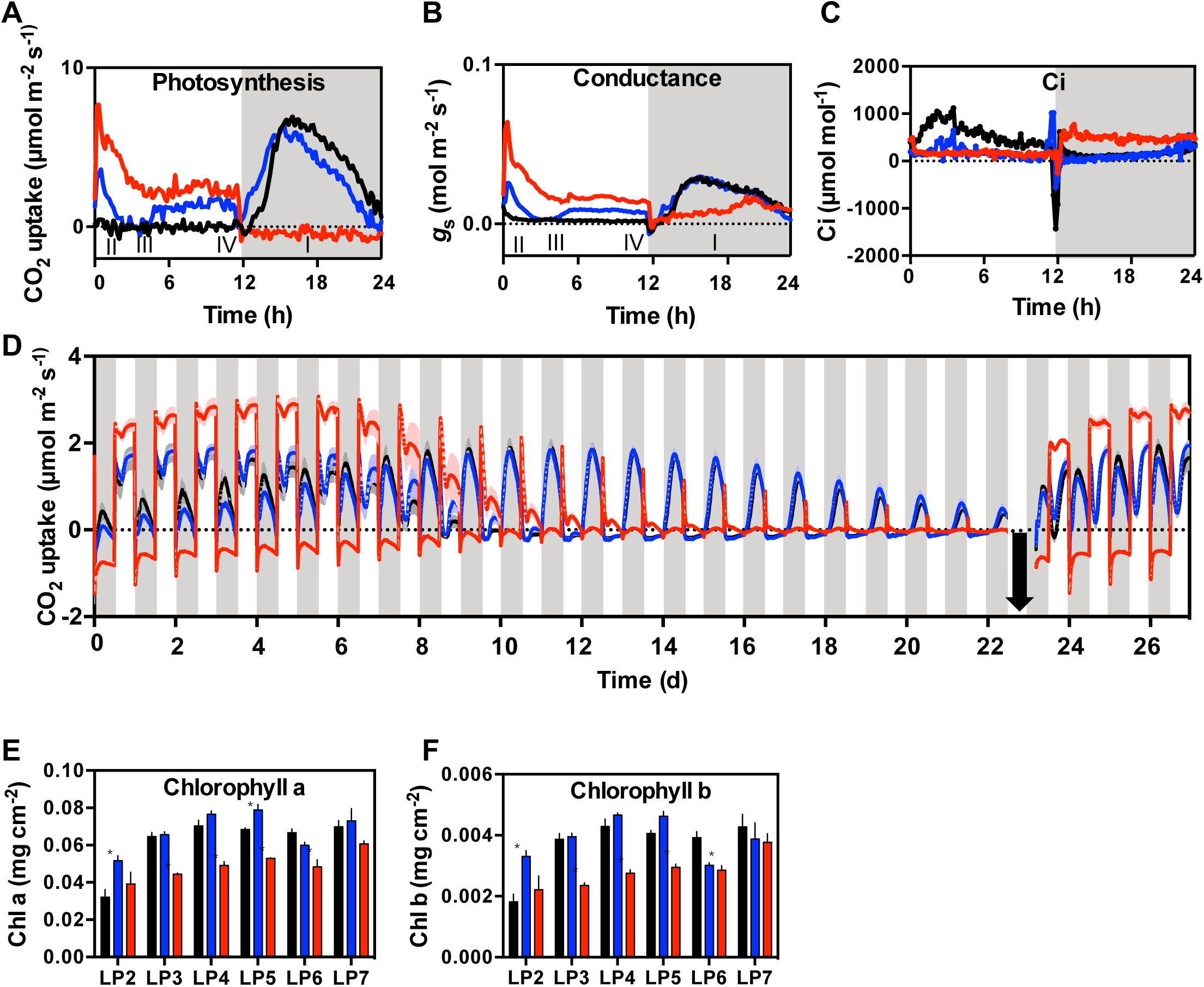
Impact of silencing *PPC1* on 24 h light/ dark gas exchange profiles under well-watered and drought-stressed conditions and chlorophyll content. (A), CO_2_ uptake profile showing the 4 phases of CAM, **(B)**, conductance profile and **(C)**, C_i_ profile for CAM leaves (leaf pair 6; LP6) using plants pre-entrained for 7-days under 12-h-light/ 12-h-dark cycles. **(D)**, Impact of drought on *K. laxiflora* wild type and transgenic lines with reduced *PPC1*. Gas exchange profile for shoots of whole young plants (9-leaf-pair stage) measured throughout 27-days. The plants were watered to full capacity on day 0 and allowed to progress into drought until day 23, when they were re-watered to full capacity (black arrow). The mean of 4 individuals is presented for each line. **(E)** and **(F)** Impact of silencing *PPC1* on chlorophyll a **(E)** and chlorophyll b **(F)** content in leaf pairs 2 through 7. In **(E)** and **(F)** the error bars represent the standard error of the mean calculated for the three biological replicates. Asterisks indicate significant difference from the wild type based on the Student’s *t* test: * **(E)**, Chlorophyll a in *rPPC1-B:* LP3 p=0.00079659, LP4 p=0.00309081, LP5 p=4.345153e-005, LP6, p=0.00330118. *rPPC1-A* in LP2, p=0.00001, LP5 p=0.02. n=3. * **(F)**, Chlorophyll b in *rPPC1-B:* LP3 p=0.00150698, LP4 p=0.00376018, LP5 p=0.00056023, LP6, p=0.0074742, n=3. *rPPC1-A* in LP2, p=0.0003. Black data are for the wild type, blue data *rPPC1-A,* and red data *rPPC1-B*.

The four phases of CAM (Osmond, 1978) are indicated on the CO_2_ exchange graph in Figure 4A. In the light period (phases II through IV), wild type leaves fixed negligible atmospheric CO_2_, whereas *rPPC1-B* fixed a total of 265 µmol m^−2^, and *rPPC1-A* fixed 89 µmol m^−2^ (Figure 4A). In the dark period (phase I), wild type fixed 297 µmol CO_2_ m^−2^ and *rPPC1-A* 226 µmol CO_2_ m^−2^, but *rPPC1-B* respired 35 µmol CO_2_ m^−2^ (Figure 4A). Loss of nocturnal CO_2_ fixation in *rPPC1-B* (Figure 4A) correlated with the lack of the CAM-associated PPC1 (Figure 1 and 2). LP6 of *rPPC1-A* fixed 24 % less nocturnal CO_2_ than wild type, but, in contrast to wild type, continued some light CO_2_ capture (Figure 4A).

Apart from opening briefly for phase II, just after lights-on, the wild type closed its stomata in the light and opened them throughout the dark, when g_s_ tracked CO_2_ uptake (Figure 4B). In *rPPC1-B*, stomata stayed open in the light, closed briefly at dusk, and opened slightly throughout the dark, with a small peak prior to dawn (Figure 4B). The dark conductance of *rPPC1-B* correlated with the release of respiratory CO_2_ (Figure 4A). The internal partial pressure of CO_2_ inside the leaf (C_i_) was highest for the wild type in the light period, when stomata were closed, whereas in *rPPC1-B,* C_i_ peaked during the dark period, when stomata were slightly open, but the leaves failed to fix atmospheric CO_2_, and respiratory CO_2_ escaped (Figure 4C).

### Impact of Drought on CO_2_ uptake in Plants lacking *PPC1*

To test the importance of *PPC1* for carbon assimilation during drought, CO_2_ uptake was measured continuously in whole, young plants (9-leaf-pairs) over 22 days of drought, followed by re-watering (Figure 4D). CAM develops with leaf age in *K. laxiflora* (Supplemental Figure S4). The leaves gradually reduce light period CO_2_ fixation and increase nocturnal CO_2_ fixation as they develop (Supplemental Figure S4). A young wild type plant with 9-leaf-pairs thus includes young leaves fixing CO_2_ mainly in the light via the C_3_ pathway, and older leaves fixing the majority of their atmospheric CO_2_ in the dark via PPC and CAM.

When W-W on day 1, wild type, *rPPC1-A* and *rPPC1-B* fixed, respectively, 7 %, 6 % and −26 % of their CO_2_ during the dark, and 93 %, 94 % and 126 % during the light period (Table 1). This indicated that, in well-watered conditions, the young leaves of these young plants performed the majority of the 24 h CO_2_ uptake (Table 1). On day 1, wild type, *rPPC1-*A and *rPPC1–B* fixed 1093, 1008 and 1078 µmol of atmospheric CO_2_ m^−2^ in total over 24 h (Table 1). After 7 days without watering, 24 h CO_2_ fixation was 1345, 1367 and 1506 µmol m^−2^. It should be noted that total leaf area was measured at the end of the experiment, so leaf growth and expansion during the experiment could not be accounted for.

**Table 1.** Analysis of the uptake of CO_2_ in wild type (WT), rPPC1-A and rPPC1-B during 22-d of progressive drought-stress and for 4 days following re-watering.

|  | Night<br>1 | Day<br>1 | Total<br>diel 1 | Night<br>7 | Day<br>7 | Total<br>diel 7 | Night<br>13 | Day<br>13 | Total<br>diel 13 | Night<br>22 | Day<br>22 | Total<br>diel 22 | Night<br>27 | Day<br>27 | Total<br>diel 27 |
| --- | --- | --- | --- | --- | --- | --- | --- | --- | --- | --- | --- | --- | --- | --- | --- |
| WT (% of diel CO<sub>2</sub> uptake) | 7 | 93 | 100 | 52 | 48 | 100 | 111 | -11 | 100 | 133 | -33 | 100 | 56 | 44 | 100 |
| WT CO<sub>2</sub> uptake (μmol m<sup>-2</sup>) | 78 | 1015 | 1093 | 698 | 647 | 1345 | 908 | -113 | 795 | 152 | -76 | 76 | 931 | 735 | 1666 |
| *rPPC1-A* (% of diel CO<sub>2</sub> uptake) | 6 | 94 | 100 | 40 | 60 | 100 | 112 | -12 | 100 | 128 | -28 | 100 | 46 | 54 | 100 |
| *rPPC1-A* CO<sub>2</sub> uptake (μmol m<sup>-2</sup>) | 73 | 1081 | 1008 | 553 | 814 | 1367 | 922 | 120 | 802 | 209 | -81 | 130 | 768 | 889 | 1657 |
| *rPPC1-B* (% of diel CO<sub>2</sub> uptake) | -26 | 126 | 100 | -14 | 114 | 100 | -6 | 106 | 100 | -27 | -73 | 100 | -17 | 117 | 100 |
| *rPPC1-B* CO<sub>2</sub> uptake (μmol m<sup>-2</sup>) | -575 | 1653 | 1078 | -254 | 1759 | 1506 | -17 | 161 | 145 | -8 | -22 | -30 | -385 | 1825 | 1440 |

Table Footnote: Data are presented for various days (1, 7, 13, 22 and 27) throughout the 27-day drought-stress and re-watering experiment. Results expressed as percentages were calculated separately for the 12-h-light and 12-h-dark periods by determining the percentage of the total diel/ 24 h CO<sub>2</sub> uptake that occurred in that period. The actual CO<sub>2</sub> uptake over the 12-h-light and 12-h-dark period is also presented for each line.

After 7 days without water, there was a substantial increase in nocturnal CO_2_ uptake in wild type and *rPPC1-A* relative to day 1 (52 % and 40 % of CO_2_ fixation occurred in the dark, respectively; Table 1). *rPPC1-B* respired less (−14 %) after 7 days of drought relative to the −26 % dark respired CO_2_ on day 1 (Table 1). After 13 days of drought, wild type, *rPPC1-A* and *rPPC1-B* fixed, respectively, 795, 802 and 145 µmol CO_2_ m^−2^ over the 24 h cycle (Table 1). Thus, plants performing CAM (wild type and *rPPC1-A*) were able to fix over 5-fold more CO_2_ after 13 days of drought compared to *rPPC1-B*. Furthermore, after 22 days of drought, wild type, *rPPC1-A* and *rPPC1-B* fixed 76, 130 and −30 µmol m^−2^ during the 24 h light/ dark cycle.

On day 23 without water, the drought-stressed plants were rewatered (see photos in Supplemental Figure S3). After re-watering, CO_2_ fixation increased rapidly for all plants (Figure 4D). In addition, wild type and *rPPC1-A,* displayed pronounced phase III of CAM after re-watering (Figure 4D). Following soil rehydration, the wild type fixed 931 µmol m^−2^ CO_2_ in the dark, compared to 768 µmol m^−2^ for *rPPC1-A*, whereas prior to drought *rPPC1-A* fixed more atmospheric CO_2_ (Table 1). *rPPC1-B* fixed CO_2_ throughout the light period following re-watering, and it also resumed respiratory CO_2_ loss throughout the dark (Figure 4D).

As there was an increase in C_3_ photosynthesis in *rPPC1-B*, chlorophyll a and b were assayed for LP2 through LP7 (Figure 4E and 4F). Line *rPPC1-A* contained significantly more chlorophyll a and b than wild type in LP2, and *rPPC1-B* contained significantly less chlorophyll a and b than the wild type in LP3 to LP6 (Figure 4E and 4F).

### Characterisation of CAM Gene Transcript Abundance in *rPPC1* Lines

Having established that *rPPC1-B* lacked nocturnal CO_2_ fixation (Figure 4A), it was important to investigate the temporal regulation of other CAM-associated genes in the *rPPC1* RNAi lines. The transcript abundance of CAM genes in CAM leaves (LP6) was investigated using samples collected every 4 h over a 12-h-light/ 12-h-dark cycle (Figure 5). *PPDK* was unchanged relative to the wild type levels in either of the *rPPC1* lines (Figure 5A), but its regulator *PPDK-RP* was up-regulated in line *rPPC1-B* (Figure 5B), consistent with the continuous phosphorylation and inactivation of PPDK (Figure 2E and 2F). *β-NAD-ME* was only slightly different from the wild type in the *rPPC1* lines, but was lower in *rPPC1-B* at 22:00 (Figure 5C).

**Figure 5.**
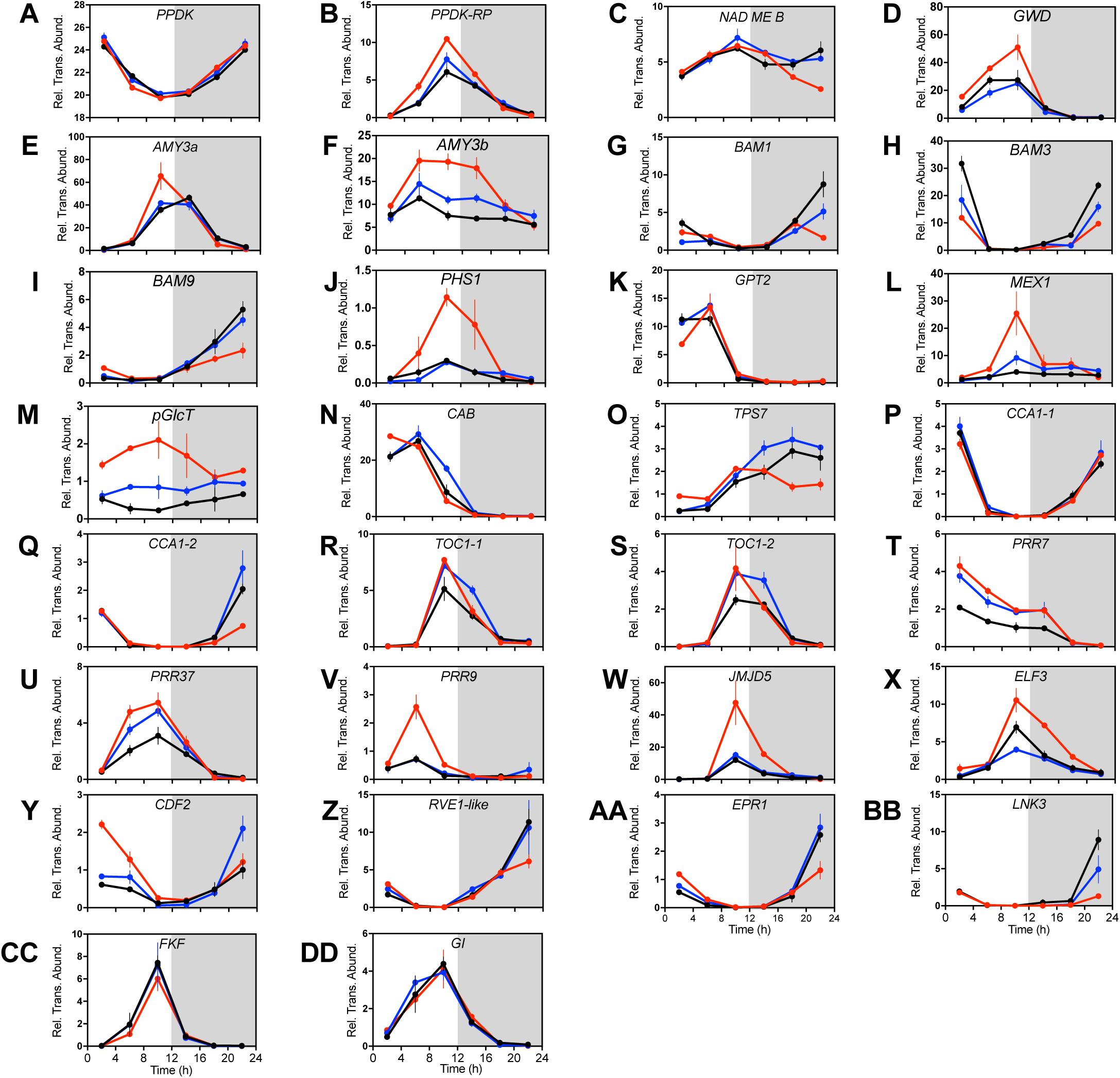
Impact of the Loss of *PPC1* on the Light/Dark Regulation of the Transcript Abundance of CAM- and Central Circadian Clock-Associated Genes in RNAi Lines *rPPC1-A* and *rPPC1-A*. Gene transcript abundance was measured using RT-qPCR for target genes: **(A)**, *PPDK*; **(B)**, *PPDK-RP*; **(C)**, β-*NAD-ME*; **(D)**, *GWD;* (E)*, AMY3a;* (F)*, AMY3b;* (G)*, BAM1;* (H)*, BAM3;* (I)*, BAM9;* (J)*, PHS1;* (K)*, GPT2;* (L)*, MEX1;* (M), *pGlcT*; **(N)**, *CAB*; **(O)**, *TPS7*; **(P)**, *CCA1-1;* (Q)*, CCA1-2;* (R), *TOC1-1;* (S)*, TOC1-2*; **(T)**, *PRR7*; **(U)**, *PRR37*; **(V)**, *PRR9*; **(W)**, *JMJ30*/ *JMJD5*; **(X)**, *ELF3;* (Y), *CDF2;* (Z), *RVE1-like;* (AA), *EPR1;* (BB), *LNK3;* (CC), *FKF1* and **(DD)**, *GI*. Mature leaves (leaf pair 6; LP6) were sampled every 4 h across the 12-h-light/ 12-h-dark cycle. A thioesterase/thiol ester dehydrase-isomerase superfamily gene (*TEDI*) was amplified from the same cDNAs as a reference gene. Gene transcript abundance data represents the mean of 3 technical replicates for biological triplicates and was normalized to loading control gene *(TEDI)*; error bars represent the standard error. In all cases, plants were entrained under 12-h-light/ 12-h-dark cycles for 7 days prior to sampling. Black data are for the wild type, blue data *rPPC1-A,* and red data *rPPC1-B*.

In light of the marked reduction in starch to half the wild type level in *rPPC1-B* (Figure 3B), transcripts associated with starch turnover were also measured. In *rPPC1-B,* dusk-phased starch breakdown-associated genes (*GWD, AMY3a and 3b, PHS1;* Figure 5D to 5F and 5J), and sugar transporters (*MEX1, pGlcT;* Figure 5L and 5M) were up-regulated, whereas dawn-phased starch breakdown genes (*BAM1, BAM3, BAM9*; Figure 5G to 5I) and sugar transporters (*GPT2;* Figure 5K) were down-regulated in comparison to the wild type.

Finally, a *CHLOROPHYLL A/B BINDING PROTEIN* (*CAB1*) gene was up-regulated in *rPPC1-B* at dawn (Figure 5N), whereas a potential sucrose sensor connecting growth and development to metabolic status, *TREHALOSE 6-PHOSPHATE SYNTHASE7* (*TPS7*) (Schluepmann et al., 2003), was down-regulated relative to the wild type at 6 and 10 h into the 12-h-dark period (Figure 5O).

### Characterisation of Diel Regulation of Circadian Clock Genes in *rPPC1* Lines

Recent studies using *PPCK1* RNAi lines of *Kalanchoë* reported that reduced sucrose 2 h after dawn correlated with perturbation of the central circadian clock (Boxall et al., 2017). Of the two *CIRCADIAN CLOCK ASSOCIATED1* (*CCA1)* genes in *K. laxiflora*, only *CCA1-2* was down-regulated in *rPPC1-B* (Figure 5P and 5Q), whereas both *TIMING OF CHLOROPHYLL A/B BINDING PROTEIN1* (*TOC1)* genes were up-regulated in both *rPPC1-A* and *rPPC1-B* (Figure 5R and 5S). Two other pseudo-response regulators (*PRRs*) related to *TOC1*, namely *PRR7* and *PRR3/7,* were up-regulated in both *rPPC1-A* and *rPPC1-B* (Figure 5T and 5U), and *PRR9* was induced almost 5-fold in the middle of the light period, specifically in line *rPPC1-B* (Figure 5V).

Other clock associated genes, including *JUMONJI DOMAIN CONTAINING30/5* (*JMJ30/ JMJD5*), *EARLY FLOWERING3* (*ELF3*) and *CYCLING DOF-FACTOR2* (*CDF2*) were also up-regulated in line *rPPC1-B* (Figure 5W to 5Y), whereas the single MYB-repeat transcription factors *REVEILLE1-like* (*RVE1-like*) and *EARLY-PHYTOCHROME-RESPONSIVE1* (*EPR1*), and *LIGHT NIGHT-INDUCIBLE AND CLOCK-REGULATED3-like* (*LNK3-like*) were down-regulated (Figure 5Z to 5BB). Finally, *CCA1-1, FLAVIN-BINDING KELCH REPEAT F-BOX PROTEIN1* (*FKF1*) and *GIGANTEA* (*GI*) levels were consistent between all three lines (Figure 5O, 5CC and 5DD).

### Gas Exchange Characteristics of *rPPC1* Lines under Circadian Free Running Conditions

Under constant light and temperature (LL) free-running conditions, detached wild type CAM leaves (LP6) of *K. laxiflora* displayed a robust circadian rhythm of CO_2_ uptake with a period of approximately 20 h (Figure 6A). This rhythm was entirely consistent with CAM rhythms reported previously for *K. fedtschenkoi* and *K. daigremontiana* (Lüttge and Ball, 1978; Anderson and Wilkins, 1989). The rhythm dampened rapidly in line *rPPC1-B* (Figure 6A), whereas *rPPC1-A* maintained a rhythm that was very similar to that of the wild type (Figure 6A).

**Figure 6.**
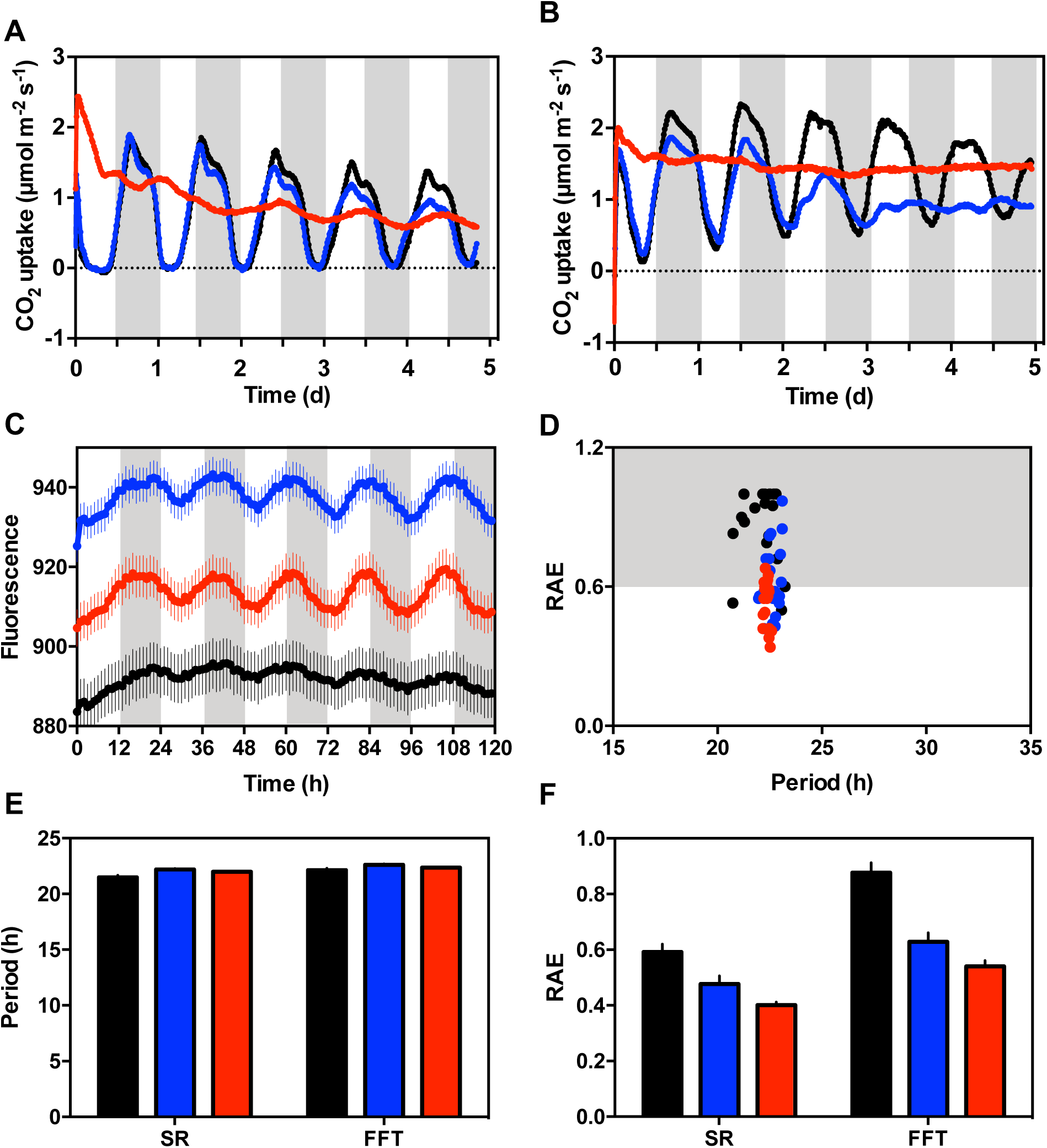
Effects of silencing *PPC1* on CAM CO_2_ exchange rhythms measured under constant light and temperature (LL) conditions. **(A)**, Gas exchange profile for detached CAM leaves (leaf pair 6; LP6) was measured using leaves entrained under a 12-h-light/ 12-h-dark cycle followed by release into LL constant conditions (100 µmol m^−2^ s^−1^ at 15 °C). **(B)**, Gas exchange profile for well-watered whole young plants (9-leaf-pairs stage) using plants entrained under 12-h-light/ 12-h-dark cycles followed by release into constant LL conditions (100 µmol m^−2^ s^−1^ at 15 °C). The data represents the mean CO_2_ uptake of 3 individual plants. **(C)**, Delayed Fluorescence (DF) circadian rhythms became more robust in lines *rPPC1-A* and *rPPC1-B*. Plants were entrained under 12-h-light/12-h-dark cycles before being transferred to constant red/blue light (35 µmol m^2^s^−1^) under the CCD imaging camera system. DF was assayed with a 1-h time resolution for 120 h. The plots represent normalized averages for DF measured for 21 leaf discs sampled from 3 biological replicates of LP6 from each line. Error bars indicate SE of the mean calculated from three biological replicates. **(D)**, Relative Amplitude Error (RAE) plot for the DF rhythms. **(E)**, Mean period length plot, and **(F)**, Mean RAE plot; the plotted values were calculated using the Biodare package for circadian rhythm analysis. FFT = fast Fourier transform-nonlinear least squares analysis, or SR = spectral resampling analysis. Black data are for the wild type, blue data *rPPC1-A,* and red data *rPPC1-B*.

When LL CO_2_ uptake was measured for well-watered whole plants, *rPPC1-B* fixed more CO_2_ than the wild type and line *rPPC1-A*; 11269, 7414, 8383 µmol CO_2_ m^−2^, respectively (Figure 6B). Wild type maintained robust oscillations of CO_2_ exchange under LL conditions, whereas *rPPC1-A* dampened to arrhythmia after 3 days, and *rPPC1-B* was arrythmic (Figure 6B).

### Delayed Fluorescence Rhythms were Amplified in *rPPC1-B* Despite the Loss of the CAM-Associated CO_2_ Uptake Rhythm

Delayed fluorescence (DF) is rhythmic in *Kalanchoë* and provides a measure of a chloroplast-derived clock-output that can be used for statistical analysis of circadian period, robustness and accuracy (Gould et al., 2009; Boxall et al., 2017). DF was measured under LL (Figure 6C), and analysed to calculate rhythm statistics using Biodare (Moore et al., 2014; Zielinski et al., 2014).

Wild type DF oscillations were very similar to those reported previously for *K. fedtschenkoi* (Figure 6C; Gould et al., 2009; Boxall et al., 2017). *rPPC1-A* and *rPPC1-B* had more robust oscillations (Figure 6C). The rhythm amplitude increased slightly with time in *rPPC1-A* and *rPPC1-B*, but remained relatively constant in the wild type. The relative amplitude error (RAE) plot showed a wider spread of period for the wild type than for the *rPPC1* lines (Figure 6D). Mean periods were between 21.5 h and 22.1 h when calculated using spectral resampling or fast Fourier transform (nonlinear least squares) methods, respectively (Figure 6E). Average periods were similar between wild type and the *rPPC1* lines. A lower mean RAE was calculated for *rPPC1-A* and *rPPC1-B* compared to the wild type (Figure 6F), supporting statistically the visibly robust and high-amplitude DF rhythm in the *rPPC1* lines (Figure 6C).

### Rhythm Characteristics of Core Circadian Clock and Clock-Controlled Genes

Having established that circadian control of CO_2_ fixation was dampened under LL, and that the circadian control of DF was enhanced under LL in plants lacking *PPC1* (Figure 6A to 6F), it was important to investigate the regulation circadian clock-controlled genes in the *rPPC1* lines under LL (Figure 7).

**Figure 7.**
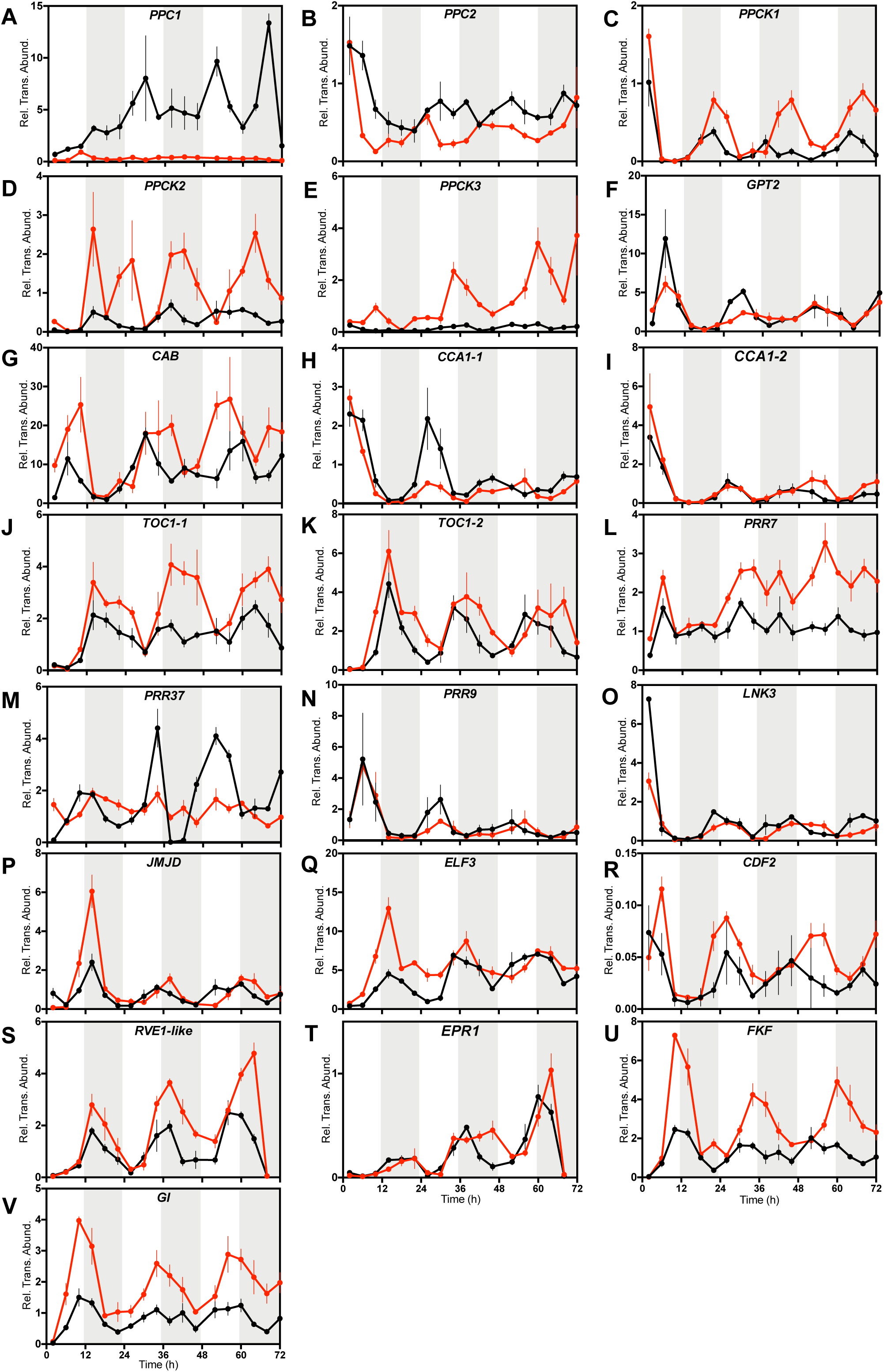
Impact of the Loss of *PPC1* activity on Circadian Clock Controlled Gene Transcript Abundance during Constant Light and Temperature (LL) Free Running Conditions. Mature leaves (leaf pair 6), which performed CAM in the wild type, were sampled every 4 h under constant conditions (100 µmol m^−2^ s^−1^ at 15 °C) for wild type *and rPPC1-B.* RNA was isolated and used for real-time RT-qPCR. A thioesterase/thiol ester dehydrase-isomerase superfamily gene (*TEDI*) was amplified as a reference gene from the same cDNAs. Gene transcript abundance data represents the mean of 3 technical replicates for biological triplicates and was normalized to loading control gene *(TEDI)*; error bars represent the standard error. In all cases, plants were entrained under 12-h-light/ 12-h-dark cycles prior to release into LL free-running conditions. Black data are for the wild type and red data *rPPC1-B.* (A), Circadian rhythm of *PPC1* transcript abundance under constant LL conditions (100 µmol m^−2^ s^−1^ at 15 °C) for wild type and *rPPC1-B*. **(B)**, *PPC2;* (C), *PPCK1;* (D), *PPCK2;* (E), *PPCK3*; **(F)**, *GPT2;* (G), *CAB*; **(H)**, *CCA1-1;* (I), *CCA1-2;* (J), *TOC1-1;* (K), *TOC1-2;* (L), *PRR7;* (M), *PRR37*; **(N)**, *PRR9*; **(O)**, *LNK3-like;* (P), *JMJ30/JMJD5;* (Q), *ELF3;* (R), *CDF2;* (S), *RVE1*; **(T)**, *EPR1*; **(U)**, *FKF1* and **(V)**, *GI*.

Wild type displayed a rhythm in the transcript abundance of *PPC1* that was absent in *rPPC1-B* (Figure 7A)*. PPC2* was rhythmic in wild type and *rPPC1-B*, but its abundance was lower in *rPPC1-B* (Figure 7B). The *PPCK1* rhythm was of greater amplitude in *rPPC1-B*, and the daily transcript peaks occurred 4 to 8 h later than the wild type after the first 24 h of LL (Figure 7C). *PPCK2* and *PPCK3* were induced in line *rPPC1-B* and oscillated with higher amplitude (Figure 7D and 7E). *PPCK2* peaked after the wild type on the second and third 24 h cycles under LL (Figure 7D and 7E), which was consistent with the induction detected under light/ dark cycles (Figure 1F and 1G). *GPT2*, involved in the transport of G6P across the chloroplast membrane, was down-regulated and had lower amplitude in *rPPC1-B* (Figure 7F), whereas *CAB1* was induced and oscillated robustly (Figure 7G).

In *rPPC1-B*, core clock gene *CCA1-1* was down-regulated, phase delayed and had lower amplitude than the wild type (Figure 7H), whereas *CCA1-2, TOC1-1* and *TOC1-2* were all up-regulated and phase delayed relative to the wild type for the latter two peaks of LL (Figure 7I, 7J and 7K). *PRR7* was up-regulated and more robustly rhythmic in *rPPC1-B* (Figure 7L), whereas the rhythms of *PRR3/7* and *PRR9* dampened in *rPPC1-B* (Figure 7M and 7N). Finally, *JMJ30/ JMJD5*, *ELF3*, *CDF2*, *RVE1-like*, *EPR1*, *FKF1* and *GI* were up-regulated in *rPPC1-B*, displaying higher amplitude, phase delays and/ or period lengthening during the LL time course (Figure 7P to 7V), whereas *LNK3-like* was down-regulated but remained rhythmic with a lengthening period (Figure 7O).

### Diel Regulation of GC-Signalling and Ion Channel Genes in GC-Enriched Epidermal Peels of *rPPC1-B*

*PHOTOTROPIN1* (*PHOT1*) encodes a protein kinase that acts as a blue light (BL) photoreceptor in the signal-transduction pathway leading to BL-induced stomatal movements (Kinoshita et al., 2001). In *rPPC1-B, PHOT1* transcripts were up-regulated relative to the wild type and peaked 8 h into the 12-h-light period (Figure 8A). *CRYPTOCHROME2* (*CRY2*) is a photoreceptor that regulates BL responses, including the entrainment of endogenous circadian rhythms (Somers et al., 1998), and stomatal conductance via an indirect effect on ABA levels (Boccalandro et al., 2012). In *rPPC1-B* epidermal peels*, CRY2* transcript levels were up-regulated and peaked at dusk, whereas *CRY2* peaked 8 h into the dark in the wild type (Figure 8B).

**Figure 8.**
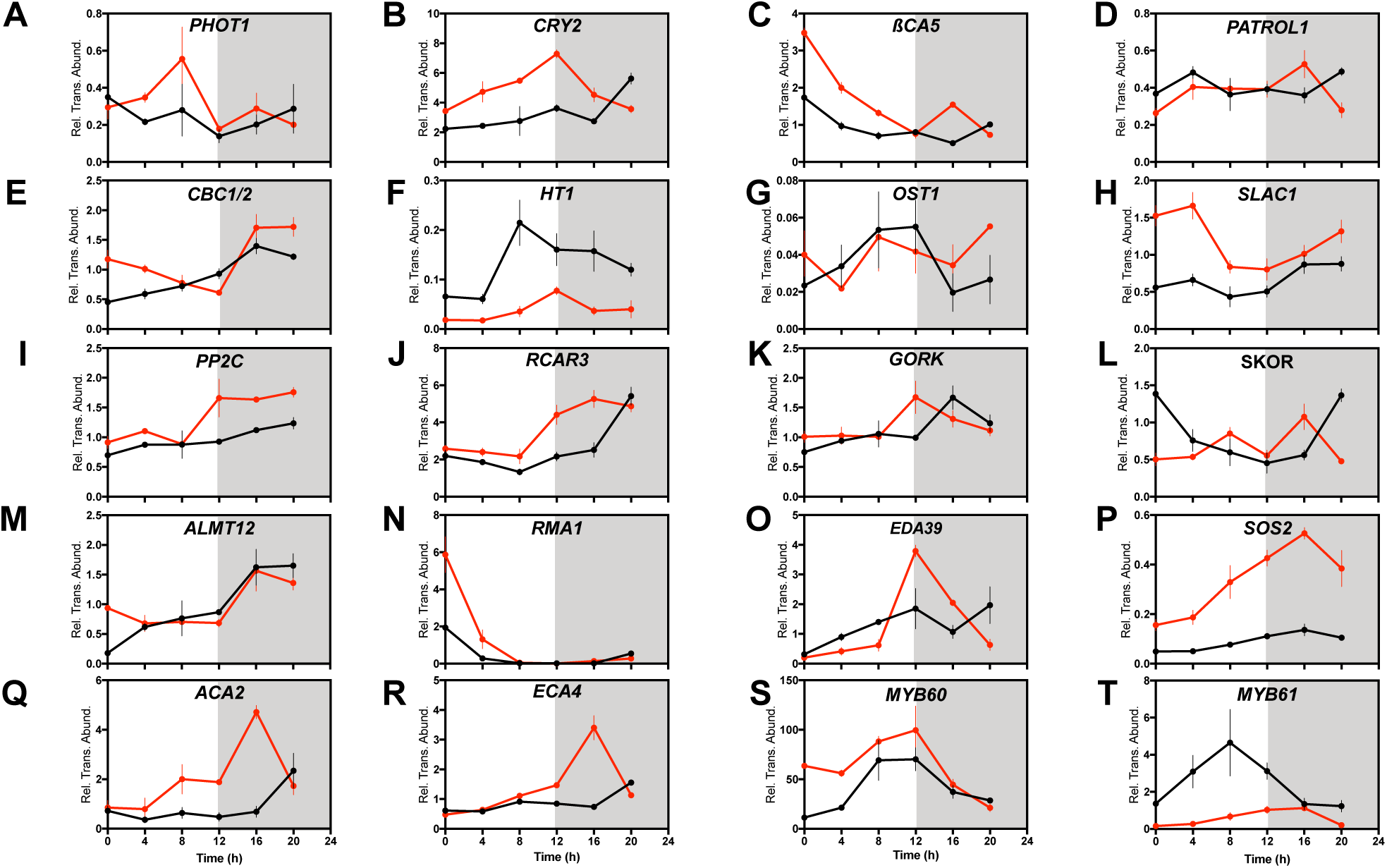
Impact of the Loss of PPC activity on the Light/ Dark Regulation of the Transcript Abundance of Stomatal Genes in the Epidermis. Plants were entrained under 12-h-light/ 12-h-dark for 7 days prior to sampling. Epidermal peel samples were separated from leaf pairs 6, 7 and 8, with samples collected every 4h starting at 02:00, 2 h into the 12-h-light period. Each biological sample represents a pool of epidermal peels taken from 6 leaf pair 6 leaves from 3 clonal stems of each line. Each peel was frozen in liquid nitrogen immediately after it was taken and pooled later. RNA was isolated and was used in RT-qPCR. A thioesterase/thiol ester dehydrase-isomerase superfamily gene (*TEDI*) was amplified as a reference gene from the same cDNAs. Gene transcript abundance data represent the mean of 3 technical replicates for biological triplicates, and was normalized to the reference gene *(TEDI)*; error bars represent the standard error. In all cases, plants were entrained under 12-h-light/ 12-h-dark cycles prior to release into LL free-running conditions. Black data are for the wild type and red data *rPPC1-B.* (A), *PHOT1*; **(B)**, *CRY2*; **(C)**, *βCA5*; **(D)**, *PATROL1*; **(E)**, *CBC1/2*; **(F)**, *HT1*; **(G)**, *OST1*; **(H)**, *SLAC1*; **(I)**, *PP2C*; **(J)**, *RCAR3*; **(K)**, *GORK*; **(L)**, *SKOR*; **(M)**, *ALMT12*; **(N)**, *RMA1*; **(O)**, *EDA39*; **(P)**, *SOS2*; **(Q)**, *ACA2*; **(R)**, *ECA4*; **(S)**, *MYB60* and **(T)**, *MYB61*.

In Arabidopsis, GC localised *β-CARBONIC ANHYDRASE1 (CA1) and CA4* are involved in CO_2_ sensing in GCs, with the *ca1ca4* mutant displaying impaired stomatal control in response to CO_2_ (Hu et al., 2010). In *rPPC1-B,* a *β-*carbonic anhydrase gene was much more strongly induced at dawn relative to the wild type (Figure 8C). *PATROL1* controls the tethering of the proton ATPase AHA1 to the plasma membrane, and is essential for opening in response to low CO_2_ and light (Hashimoto-Sugimoto et al., 2013). The cycle of *PATROL1* in epidermal peels was slightly different between *rPPC1-B* and wild type, with *rPPC1-B* peaking 4 h into the dark, but the wild type peak at 4 h into the light (Figure 8D).

*CONVERGENCE OF BLUE LIGHT AND CO2 1/2* (*CBC1/2*) stimulate stomatal opening by inhibiting S-type anion channels in response to both BL and low CO2 and are known to connect BL signals perceived by PHOT1 with the protein kinase *HIGH LEAF TEMPERATURE1* (*HT1*) (Hiyama et al., 2017). Both proteins interact with and are phosphorylated by HT1. In *rPPC1-B, CBC1/2* was up-regulated compared to the wild type both in the second half of the dark period and at dawn (Figure 8E). *HT1* acts as a negative regulator of high CO_2_ induced stomatal closure (Hashimoto et al., 2006). *HT1* transcripts were repressed throughout the light/ dark cycle relative to the wild type in *rPPC1-B* (Figure 8F). *OPEN STOMATA1 (OST1*), a further protein kinase that acts downstream of *HT1* (Imes et al., 2013), was up-regulated in *rPPC1-B* both at 8h into the dark, and at dawn, the time of the daily trough in the wild type (Figure 8G).

Genes involved in ion transport across the plasma membrane and tonoplast play a crucial role in the cell turgor pressure changes that drive stomatal movements (Jezek and Blatt, 2017). *SLOW ACTIVATED ANION CHANNEL1* (*SLAC1*) is a key player in the closure of stomata in response to high CO_2_ concentration (Hedrich and Geiger, 2017). *SLAC1* is regulated by ABA signalling that requires de-phosphorylation steps catalysed by protein phosphatase type 2C (*PP2C*). In *rPPC1-B, SLAC1* was up-regulated and peaked during the light (Figure 8H), and *PP2C* was up-regulated relative to the wild type during the dark period (Figure 8I). ABA is recognized and bound by the *REGULATORY COMPONENT OF ABA RECEPTORS* (*RCAR*s)/ *PYRABACTIN RESISTANCE1 (PYR1*/*PYL)* and interacts with PP2C to stimulate ABA signalling (Ma et al., 2009; Park et al., 2009; Santiago et al., 2009). In *rPPC1-B* epidermal peels*, RCAR3* transcripts were induced spanning dusk and the first half of the dark period, and peaked at least 4 h earlier than the wild type (Figure 8J).

In *rPPC1-B,* the *GUARD CELL OUTWARD RECTIFYING K^+^ CHANNEL* (*GORK*), which is known to be involved in the regulation of stomata according to water status (Ache et al., 2000), peaked 4 h earlier than wild type (Figure 8K). *STELAR K^+^ OUTSIDE RECTIFIER* (*SKOR)* is a selective outward-rectifying potassium channel (Gaymard et al., 1998). It peaked at dawn in the wild type, whereas in *rPPC1-B*, *SKOR* transcript levels reached their daily minimum at dawn (Figure 8L). The plasma-membrane localised *ALUMINIUM ACTIVATED MALATE TRANSPORTER12* (*ALMT12*), which is involved in dark-, CO_2_-, abscisic acid- and water deficit-induced stomatal closure, was elevated relative to the wild type at dawn in *rPPC1-B* (Figure 8M). An *E3 UBIQUITIN-PROTEIN LIGASE* (*RMA1*) promotes the ubiquitination and proteasomal degradation of aquaporin PIP2:1, which is known to play a role in GC regulation (Grondin et al., 2015). *RMA1* was induced 3-fold at dawn in *rPPC1-B* compared to the wild type (Figure 8N). *EMBRYO SAC DEVELOPMENT ARREST39* (*EDA39*) is a calmodulin binding protein that promotes stomatal opening (Zhou et al., 2012). In *rPPC1-B*, *EDA39* was up-regulated relative to the wild type and peaked at dusk (Figure 8O).

*SALT OVERLY SENSITIVE2* (*SOS2*) is a Calcineurin B-Like (CBL)-interacting protein kinase involved in the regulatory pathway for intracellular Na^+^ and K^+^ homeostasis and salt tolerance (Liu et al., 2000). SOS2 has also been demonstrated to interact with and activate the vacuolar H^+^/ Ca^2+^ antiporter CAX1, thereby functioning in cellular Ca^2+^ homeostasis; an important function during stomatal opening and closing (Cheng et al., 2004). *SOS2* transcript levels were elevated relative to the wild type at all time points, and in particular, it was induced ∼5-fold at its peak 4 h into the dark period in epidermal peels of *rPPC1-B* relative to the wild type (Figure 8P). The *Ca^2+^-ATPase2* (*ACA2*) and *ENDOMEMBRANE-TYPE Ca^2+^-ATPASE4* (*ECA4*) catalyze the hydrolysis of ATP coupled with the translocation of calcium from the cytosol into the endoplasmic reticulum and/ or an endomembrane compartment (Jezek and Blatt, 2017). *ACA2* and *ECA4* were induced by between ∼4-fold to ∼8-fold in *rPPC1-B* epidermal peels relative to the wild type, particularly when they reached peak levels at 4 h into the 12-h-dark period (Figure 8Q and 8R).

Finally, transcription factor *MYB60* is involved in stomatal opening in response to light and also promotes GC deflation in response to water deficit (Cominelli et al., 2005). In *rPPC1-B, MYB60* was induced ∼5-fold relative to the wild type at dawn, although it also maintained a dusk phased peak of transcript abundance like the wild type (Figure 8S). *MYB61* functions as a transcriptional regulator of stomatal closure (Liang et al., 2005). It was down-regulated in *rPPC1-B* relative to wild type, and peaked 8 h later in *rPPC1-B,* 4 h into the dark period, whereas wild type *MYB61* peaked in the light 4 h before dusk (Figure 8T).

## DISCUSSION

### Gene Specific Silencing of the CAM Isogene *PPC1* Responsible for Dark CO_2_ Fixation in an Obligate CAM Plant

*PPC1* loss-of-function mutants of the obligate CAM species *K. laxiflora* were generated using RNAi (Figure 1 and 2). Although *PPC2* transcripts were induced almost 7-fold in the most strongly silenced line, *rPPC1-B* (Figure 1B), this did not compensate for the loss of the CAM-associated *PPC1* (Figure 1A and 2A). Total extractable PPC activity was below the level of detection in *rPPC1-B* (Figure 2E), and PPC-catalyzed dark period fixation of atmospheric CO_2_ was abolished (Figure 4A and 4D). Furthermore, transcript abundance of the PPC1 regulatory protein kinase, *PPCK1,* was reduced at the time of its circadian clock-mediated nocturnal peak (Figure 1E), and *PPCK2* and *PPCK3* were up-regulated (Figure 1F and 1G), but no phosphorylation of PPC was detected in line *rPPC1-B* (Figure 2B). This outcome was most likely due to the fact that the major substrate for PPCK1, namely the CAM-associated PPC1, had been down-regulated such that there was no substrate for any PPCK activity that resulted from translation of the available *PPCK1*, *PPCK2,* or *PPCK3* transcripts (Figure 1). This suggests that the induction of *PPCK2 and PPCK3* transcripts may have been a futile attempt to compensate for the loss of *PPC1* through attempting to enhance the nocturnal activation of any remaining PPC protein.

### Physiological Consequences of Silencing CAM Isogene *PPC1* in *K. laxiflora*

In mature leaves (LP6) and young whole plants (9-leaf-pairs stage) of *rPPC1-B*, the timing of primary CO_2_ fixation switched from the dark period to the light (Figure 4A and 4D), and the main period of stomatal conductance shifted to the light period (Figure 4B). This revealed that primary atmospheric CO_2_ fixation was occurring in the light via direct RuBisCO-mediated CO_2_ fixation into the Calvin-Benson Cycle in *rPPC1-B* (Figure 4). Consistent with the gas exchange phenotype, the amounts of malate, starch and soluble sugars were reduced in *rPPC1-B* CAM leaves (Figure 3), and this correlated with the small but significant negative impact on plant yield (Supplemental Figure S2).

Gas exchange measurements using whole plants revealed that *rPPC1-B* fixed 82% of the total amount of atmospheric CO_2_ fixed over 24 h by the wild type (Day 1, Table 1). This emphasised that well-watered wild type fixed only 18 % of their daily CO_2_ in the dark via PPC (Table 1). However, this changed dramatically during drought stress (Figure 4D). The wild type and intermediate line *rPPC1-A* increased nocturnal CO_2_ fixation throughout 22 days of drought (Figure 4D); fixing 100 % of their atmospheric CO_2_ in the dark via PPC after both 13 and 22 days of drought (Table 1). By contrast, *rPPC1-B* failed to achieve net dark period atmospheric CO_2_ fixation throughout 22 days of drought stress (Figure 4 and Table 1). Furthermore, the light-period CO_2_ fixation of *rPPC1-B* collapsed rapidly after day 7 of water-with-holding, dropping from 1759 µmoles CO_2_ m^−2^ over the 12-h-light period on day 7, to 161 µmoles m^−2^ on day 13, and net respiratory CO_2_ loss of −22 µmoles m^−2^ on day 22 (Table 1). These data demonstrate very clearly the importance of a fully functional CAM system for continued atmospheric CO_2_ fixation throughout a period of drought lasting just over 3-weeks.

Despite the inability of *rPPC1-B* to induce CAM during drought, the domed shape of its respiratory CO_2_ release throughout the 12-h-dark period (Figure 4D) indicated active refixation of respiratory CO_2_ that peaked around the middle of each 12-h-dark period. This revealed that these plants were still capable of performing a version of 24-h photosynthetic physiology approximating CAM-idling (Winter, 2019). This was further supported by the low level of malate accumulation at dawn that was achieved by *rPPC1-B* (Figure 3A), and the decline and/ or midday dip in atmospheric CO_2_ fixation in the light period (Figure 4A and 4D).

Our data also revealed that the intermediate line *rPPC1-A* was less able to adapt to drought by inducing CAM when compared to the wild type. After only 7 days of drought, the wild type had up-regulated dark period CO_2_ fixation and down-regulated light period CO_2_ fixation such that 52 % (698 µmoles m^−2^) of daily atmospheric CO_2_ fixation was occurring in the 12-h-dark period; a sharp rise from 7 % (78 µmoles m^−2^) on day one (Table 1). By contrast, line *rPPC1-A* had only progressed to performing 40 % (553 µmoles m^−2^) of its daily atmospheric CO_2_ fixation in the dark period on day 7 of drought (Table 1). This revealed that the intermediate level of silencing of *PPC1* in line *rPPC1-A* (Figure 1A, 2A and 2C) reduced the rate at which the whole plants increased the proportion of atmospheric CO_2_ fixed in the dark period, and decreased the proportion of atmospheric CO_2_ fixed in the light period, in response to drought (Figure 4D and Table 1).

Upon re-watering on day 23, wild type and *rPPC1-A* returned to performing CAM, including pronounced phase II and IV at the start and end of the light period, respectively (Figure 4D). It was noteworthy that the wild type performed more dark period CO_2_ fixation than *rPPC1-A* following re-watering (Figure 4D), suggesting that the wild type had an advantage in terms of achieving more dark CO_2_ fixation, both in the well-watered young plants on days 1 through 10 of drought (Figure 4D), and when subsequently recovering higher levels of dark CO_2_ fixation upon re-watering on day-23 (Figure 4D). Line *rPPC1-B* also bounced back to its normal, pre-drought physiology after re-watering, which was consistent with the re-watering response reported previously for a range of facultative, weak-CAM species, including *M. crystallinum*, *Portulaca oleracea*, *P. umbraticola*, *T. triangulare*, and various *Calandrinia* species (Winter and Holtum, 2014; Holtum et al., 2017; Winter, 2019).

Although the *rPPC1-B* plants fixed all of their CO_2_ in the light, especially during a pronounced phase II in the hours after dawn, the CO_2_ uptake pattern was not constant over the light period (Figure 4A). For example, in LP6 of a well-watered plant, CO_2_ fixation dropped from ∼8 µmol m^−2^ s^−1^ during phase II after dawn, to ∼2 to 3 µmol m^−2^ s^−1^ around 4 h after lights on (Figure 4A), and stomatal conductance reduced to a similar extent (Figure 4B). The observed stomatal closure could not have been due to a high malate concentration and associated internal release of CO_2_ in the mesophyll (Figure 3A). These data are consistent with the current hypothesis that the circadian clock output may drive the closure of stomata and the associated decline in atmospheric CO_2_ fixation, even when a CAM leaf has not produced malate during the preceding dark period (Von Caemmerer and Griffiths, 2009).

Overall, with respect to CAM physiology, the data presented here provide strong support for the long-held view that an ability to use CAM, and rely increasingly on CAM in response to drought stress, provides a genuine adaptive advantage in terms of prolonging net atmospheric CO_2_ fixation during drought progression (Figure 4D) (Kluge and Fischer, 1967; Osmond, 1978). The data demonstrated that the wild type and line *rPPC1-A* achieved, respectively, net atmospheric CO_2_ fixation of 18,568 and 19,071 µmoles m^−2^ over the 22-day drought progression, whereas line *rPPC1-B* only achieved 13,983 µmoles m^−2^ net atmospheric CO_2_ fixation (Supplemental Table S2). Thus, the wild type, with its fully functional CAM system, was able to fix 33 % greater more CO_2_ over the entire drought treatment period.

### Consequences of the loss of *PPC1* for the regulation of CAM-associated genes under LD cycles

Transcripts for the PPDK post-translational regulatory protein, *PPDK-RP,* plus those encoding the β-subunit of *NAD-ME*, which performs malate decarboxylation in the light and generates the substrate for PPDK (Dever et al., 2015), were both perturbed in *rPPC1-B* (Figure 5B and 5C). The induction of *PPDK-RP* correlated with the continued phosphorylation of PPDK in the light period in *rPPC1-B* (Figure 2F), and the repression of *β-NAD-ME* transcripts was consistent with a reduced demand for malate decarboxylation in the light period in the absence of substantial malate accumulation at dawn (Figure 3A).

Starch, sucrose, glucose and fructose levels were also reduced relative to the wild type in *rPPC1-B* (Figure 3B to 3E). Starch storing CAM species are believed to employ the phosphorylytic pathway for starch breakdown, and export mainly G6P from their chloroplasts in the dark (Neuhaus and Schulte, 1996; Borland et al., 2009; Borland et al., 2016), whereas C_3_ species such as *A. thaliana* are known to use the amylolytic pathway for nocturnal starch degradation and export maltose and glucose (Zeeman et al., 2010; Weise et al., 2011). Temporal profiling of starch breakdown-associated transcripts revealed some marked perturbations in transcript abundance and timing (Figure 5D to 5M). *GWD*, *AMY3a*, *AMY3b*, *PHS1*, *MEX1* and *pGlcT* were all induced in *rPPC1-B* relative to the wild type, whereas *BAM1*, *BAM3*, *BAM9* and *GPT2* were all repressed at their respective peaks in *rPPC1-B* (Figure 5). The induction of both *AMY3* isogenes and *PHS1* supports the proposal that the *rPPC1-B* line redirected more of its starch degradation through starch phosphorylase. However, this would require export of G6P via GPT, and yet *GPT2* transcript levels were repressed (Figure 5K). In addition, *GWD* was also induced (Figure 5D), suggesting that the mesophyll cells also increased starch breakdown via GWD and β-amylase, although all measured *BAM* transcripts were repressed (Figure 5G, 5H and 5I).

As *MEX1* and *pGlcT* were induced with a peak phased to dusk (Figure 5L and 5M), and *GPT2* was repressed at 2 h into the light period (Figure 5K), we favour the proposal that the *rPPC1-B* line redirects more of its nocturnal starch degradation via the amylolytic route, with *MEX1* and *pGlcT* acting to export the products of starch breakdown from the chloroplast in the dark as maltose and glucose, respectively. This proposal is also consistent with the overall return of *rPPC1-B* to a more C_3_-like pathway of CO_2_ fixation and associated metabolism, especially given that the accepted C_3_ pathway of leaf starch breakdown in the dark uses amylolytic route involving the MEX1 and pGlcT transporters (Zeeman et al., 2010).

### Core clock gene regulation under LD cycles in the absence of PPC1

In CAM *Kalanchoës*, the circadian clock optimises the timing of the daily cycle of dark CO_2_ fixation via PPC, and light period malate turnover and CO_2_ refixation via RuBisCO (Hartwell, 2006; Boxall et al., 2017). In line *rPPC1-B,* some, but not all, core circadian clock-associated genes displayed altered temporal profiles of transcript abundance under LD cycles (Figure 5O to 5DD). *CCA1-1*, *FKF1* and *GI* transcript levels were remarkably consistent between all three lines (Figure 5O, 5CC and 5DD), revealing that individual genes belonging to the core and evening loops of the core oscillator (Pruneda-Paz and Kay, 2010), maintained their patterns of regulation in the absence of *PPC1* and CAM (Figure 5). However, the LD cycles of numerous other core clock-component genes and clock-associated genes were altered in line *rPPC1B* (Figure 5). It was particularly noteworthy that core clock loop components *TOC1-1* and *TOC1-2* were induced at the time of their dusk phased peak, and *CCA1-2* was repressed ∼4-fold at its peak 2 h before dawn (Figure 5P, 5Q and 5R). Morning loop components *PRR7*, *PRR3/7* and *PRR9* were all induced, but had a similar temporal pattern to the wild type (Figure 5T, 5U and 5V), whereas *RVE1-like*, *EPR1* and *LNK3* were all repressed, correlating with the down-regulation of *CCA1-2* at the same time point, 2 h before dawn (Figure 5P, 5Z, 5AA and 5BB). In addition, Evening Complex (EC) component *ELF3* was induced at the time of its peak, 2 h before dusk, and clock-associated gene *JMJ30/ JMJD5,* which encodes an histone demethylase that functions in the regulation of circadian period and temperature compensation of the core oscillator (Jones et al., 2010; Lu et al., 2011; Jones et al., 2019), was also induced 3-fold at its peak at the same time (Figure 5W and 5X). Finally, *CDF2*, a clock-output pathway component encoding a DOF family transcription factor that regulates flowering time in Arabidopsis (Fornara et al., 2009), was induced 4-fold at 2 h after dawn and peaked 4 h later than wild type in *rPPC1-B* (Figure 5Y). In Arabidopsis, CDFs are known to repress the level of the flowering time regulator CONSTANS (CO), and their level is controlled as an output from the clock through the combined interaction of FKF1 and GI; GI stabilises FKF1, which interacts with CDFs and drives their degradation, which in turn leads to the de-repression of *CO* (Fornara et al., 2009). It is therefore intriguing that *rPPC1-B* had elevated *CDF2* at 2 h after dawn, and yet the transcript levels of *FKF1* and *GI* were very similar between wild type and *rPPC1-B* (Figure 5Y, 5CC and 5DD). Furthermore, the *rPPC1-B* line does not flower constitutively under non-inductive long days, although we have not subjected the plants to inductive short days to investigate whether they flower faster than the wild type under inductive conditions.

### Impact of PPC Silencing on the CAM Circadian Rhythm of CO_2_ Fixation

Under continuous light and temperature (LL) conditions, the CAM-associated circadian rhythm of CO_2_ fixation was dampened towards almost complete arrhythmia in detached leaves and whole, young plants of *rPPC1-B,* and CO_2_ was fixed continuously (Figure 6A and 6B). The collapse of the CO_2_ circadian rhythm suggests that CO_2_ fixation by RuBisCO in *rPPC1-B* was not subject to robust and high amplitude circadian control; certainly not in comparison to the rhythm of atmospheric CO_2_ fixation during CAM in the wild type (Figure 6B). In *K. daigremontiana,* it has been suggested that RuBisCO made a large contribution to the observed rhythm of CO_2_ assimilation under LL conditions, as the level of malate did not oscillate (Wyka and Lüttge, 2003). By contrast, online carbon isotope discrimination measurements demonstrated that the LL CO_2_ rhythm of CAM leaves of *M. crystallinum* had a RuBisCO discrimination early in each subjective dark period, which then transitioned to a PPC isotope ratio later in the subjective dark (Davies and Griffiths, 2012). In *rPPC1-B* with no detectable PPC, the remaining RuBisCO-mediated rhythm displayed, at best, only a very weak rhythm of CO_2_ assimilation (Figure 6A). This suggests that the LL rhythm has only a small contribution from C_3_ carboxylation via RuBisCO in *K. laxiflora* CAM leaves, or, perhaps, that the robust, high amplitude CO_2_ rhythm of the wild type depends on interplay between high PPC activity and RuBisCO, which ceases in the absence of PPC1. Overall, the weakly rhythmic to arrhythmic pattern of CO_2_ assimilation in *rPPC1-B* under LL was very similar to that of C_3_ *M. crystallinum* (Davies and Griffiths, 2012).

The rhythm of CO_2_ fixation in LL conditions is regulated via phosphorylation of PPC by circadian clock controlled PPCK1 (Hartwell, 2006; Boxall et al., 2017). In *rPPC1-B*, *PPCK1* was slightly up-regulated and displayed a robust, high amplitude rhythm (Figure 7C). However, as *PPCK1* had no substrate PPC to phosphorylate (Figure 2B), the data presented here provide further support for the proposal that the circadian control of CAM via *PPCK1* is dependent on PPC phosphorylation, and not in some way linked directly to the circadian oscillations of *PPCK1* transcript and activity levels (Boxall et al., 2017).

Furthermore, in addition to the robust rhythm of *PPCK1*, transcript levels of *PPCK2* and *PPCK3* were also induced and became dramatically more rhythmic under LL (Figure 7). The direct functions of *PPCK2* and *PPCK3* are currently unknown, but they may be involved in phosphorylating other PPCs in different tissues or cell types, with one possible location being the GCs, as supported by the epidermal peels RT-qPCR data in Figure 8. Overall, the induction and robust rhythmicity of *PPCK2* and *PPCK3* transcripts was futile with respect to CAM circadian physiology, as they failed to drive a robust rhythm of CO_2_ assimilation under LL conditions (Figure 6A and 6B).

### Regulation of core clock genes in *rPPC1-B*

Similar to the LL CO_2_ fixation phenotype reported here for *rPPC1-B*, transgenic *K. fedtschenkoi* lines lacking *PPCK1* also lost the CO_2_ fixation rhythm, and the transcript oscillations of many core clock genes were altered (Boxall et al., 2017). However, different core clock genes were perturbed in *rPPC1-B* compared to those whose rhythmic regulation changed in the *rPPCK1* lines of *K. fedtschenkoi* (Boxall et al., 2017 cf. Figure 7). In *rPPC1-B*, *CCA1-1* transcript oscillations dampened, and both *TOC1-1* and *TOC1-2* transcripts were up-regulated and oscillated with delayed peak phase relative to the wild type (Figure 7H, 7J and 7K). By contrast, in line *rPPCK1-3*, *CCA1-1, CCA1-2* and *TOC1-2* dampened rapidly towards arrhythmia under LL free-running conditions, whereas *TOC1-1* was up-regulated and rhythmic (Boxall et al., 2017). Furthermore, whilst *PRR7* was up-regulated and rhythmic under LL in *rPPC1-B* (Figure 7), after an initial induced peak during the first 6 h of LL, the same gene dampened to arrhythmia in *rPPCK1-3* (Boxall et al., 2017). The evening phased genes *GI* and *FKF1* were induced and rhythmic under LL conditions in both *rPPC1-B* and *rPPCK1-3,* but the level of transcript induction and the rhythm amplitude were much greater in *rPPC1-B* (Figure 7U and 7V).

Overall, these differences between the rhythms of core clock genes in *rPPC1-B* and *rPPCK1-3* revealed that the clock responded very differently to the silencing of two interconnected genes that lie at the heart of nocturnal CO_2_ fixation. Elucidating the mechanistic basis for these differences should be a fruitful avenue for further investigation, especially in light of the proposed cross-talk between CAM-associated metabolites and regulation within the core clock (Boxall et al., 2017), which was further supported by the clock gene phenotypes reported here (Figure 7).

### Interactions between sugars linked to CAM and the core circadian clock

In both LD and LL conditions, *TOC1-1*, *TOC1-2*, *PRR7* and *PRR3/7* were up-regulated in *rPPC1-B*, although only at certain time points in the subjective light and dark periods in the case of *PRR3/7* under LL (Figure 5I to 5L, and 7J to 7M). In Arabidopsis, *PRR7* is required for sensing metabolic status and coordinating the clock with photosynthesis (Haydon et al., 2013). In *rPPC1-B*, the low sucrose at 2 h after dawn (Figure 3C) may be sensed via a mechanism involving *PRR7* (Boxall et al., 2017). However, in *Kalanchoë,* a related-*PRR* gene, *PRR3/7*, was the more abundant transcript and displayed a transcript peak 2 h before dusk under LD (Figure 5T). In the wild type, *PRR3/7* may function as part of a signal transduction pathway that senses metabolic status at dusk when primary CO_2_ fixation begins (Haydon et al., 2013; Boxall et al., 2017).

A further insight into the perturbation of sugar sensing and signalling in line *rPPC1-B* resulted from the discovery that the regulation of *TREHALOSE 6-PHOSPHATE SYNTHASE7* (*TPS7*) was altered in *rPPC1-B* (Figure 5O). *TPS7* is a member of the class II *TPS* genes that have both inactive TPS and TP-phosphatase (TPP) domains, but which are proposed to play a role in signalling metabolic status of the cell in Arabidopsis (Ramon et al., 2009). In *rPPC1-B*, *TPS7* was down-regulated relative to the wild type (Figure 5N). The *TPS7* ortholog in *K. fedtschenkoi* was also down-regulated in transgenic lines that had reduced CAM and less robust rhythms of CAM-associated CO_2_ fixation due to the silencing and down-regulation of either *PPCK1*, *β*-*NADME* or *PPDK* (Dever et al., 2015; Boxall et al., 2017). Thus, *TPS7* in *Kalanchoë* may function in the sensing and/ or signalling of the cellular metabolic status. However, there are currently no data relating to a potential role for *TPS7* in relation to sugar sensing leading to metabolic entrainment of the core circadian oscillator in either *Kalanchoë* or other plant species (Ramon et al., 2009).

### Robust rhythms of DF in the absence of *rPPC1-B*, but arrhythmic DF in the absence of the PPC1 circadian phospho-regulator *PPCK1*

One of the most noteworthy circadian phenotypes related to the contrast between the current findings of robust and increased amplitude DF rhythmicity in *rPPC1-A* and *rPPC1-B* (Figure 6C), and the arrhythmic DF reported previously for the *K. fedtschenkoi rPPCK1-3* line (Boxall et al., 2017). This fundamental difference provides potentially novel insights into the underpinning regulators of the DF rhythm originating from inside the chloroplasts. In terms of core clock gene regulation under both LD and LL, both *rPPC1-B* and *rPPCK1-3* displayed similar changes in the rhythms of *TOC1-1*, *GI* and *FKF1*, as all three genes were induced and more robustly rhythmic under LL. Furthermore, the oscillations of *CCA1-1*, *PRR3/7*, *PRR9*, and *LNK3-like* all displayed reduced transcript abundance and/ or rhythm dampening in both transgenic lines. However, *CCA1-2*, *TOC1-2*, *PRR7* and clock-controlled gene *CDF2*, were all induced and rhythmic uniquely in *rPPC1-B.* Thus, *CCA1-2*, *TOC1-2*, *PRR7* and *CDF2* are the most likely candidates for driving the induction and robust LL rhythmicity of DF in *rPPC1-B*. Conversely, the decline in the amount and rhythmicity of these transcripts in *rPPCK1-3* may play a role in the dampening of the DF rhythms in that transgenic line (Boxall et al., 2017).

These results also allow us to propose that robust rhythmicity of the CAM-associated CO_2_ fixation rhythm in wild type *Kalanchoë* is most likely to rely on *CCA1-1*, *PRR3/7, PRR9*, and *LNK3-like*. However, it must be stressed that the limited set of core clock and clock-associated genes profiled under LL to date in *rPPC1-B* and *rPPCK1-3* means that there are likely to be other clock-associated genes involved in driving robust CAM CO_2_ fixation rhythms, and robust DF rhythms.

### Perturbation of diel rhythms of gene transcript oscillations in GCs

A key gap in current understanding of the molecular-genetics and physiology associated with CAM centres on the cell signalling mechanisms that mediate the inverse pattern, relative to C_3_, of stomatal opening and closing (Borland et al., 2014; Males and Griffiths, 2017). It has been proposed that GCs of CAM leaves and stems respond directly to the internal supply of CO_2_. This theory has recently led several groups to use whole leaf RNA-seq datasets to investigate alterations, relative to C_3_ leaves, in the temporal phasing of known GC regulatory genes and membrane transporters (Abraham et al., 2016; Wai and VanBuren, 2018; Yin et al., 2018; Heyduk et al., 2019; Moseley et al., 2019). However, these studies did not use enriched GCs as the source of RNA, and therefore the data that were mined represented all leaf cell types, including palisade and spongy mesophyll, GCs, subsidiary cells, phloem, phloem companion cells, xylem, bundle sheath, and water storage parenchyma in Agave and pineapple. Thus, the re-scheduling of the temporal patterns of candidate GC genes in these previous datasets may have been complicated by transcripts from the same genes that were functional in other cell types of the leaves.

Separated epidermal peels from *Kalanchoë* CAM leaves are enriched for intact guard cells. We leveraged this feature in order to compare the temporal pattern of transcript regulation between GC-enriched epidermal peels of the wild type and *rPPC1-B* (Figure 8). A wide range of genes known to be involved in stomatal opening and closing displayed alterations in both transcript abundance, and/ or the timing of the daily transcript peak (Figure 8A to 8U). In particular, the temporal re-phasing and/ or down-regulation/ up-regulation of GC transcripts including *HT1*, *OST1*, *SLAC1*, *RCAR3*, *GORK*, *SKOR*, *ALMT12*, *RMA1*, *EDA39*, *SOS2*, *ACA2*, *ECA4*, and *MYB*s *60*, and *61,* revealed that the temporal control of the regulatory circuits that mediate the opening and closing of stomata was altered in *rPPC1-B* (Figure 8). It must however be emphasised that transcript changes alone are often difficult to interpret as they do not provide direct evidence concerning the encoded protein’s abundance and activity. Despite this caveat, the GC transcript abundance alterations in *rPPC1-B* (Figure 8) correlated with the 12-h-shift in stomatal opening to the light period in *rPPC1-B* at the level of whole leaf physiology (Figure 4B). Taken together, these results therefore support the theory that the measured changes in transcript abundance and temporal patterns for a range of guard cell regulatory genes were connected to the measured changes in stomatal opening and closing.

Thus, the up-regulation of *HT1*, *SKOR* and *MYB61* in the wild type relative to *rPPC1-B*, and the down-regulation of a wide range of genes, including *SLAC1*, *PP2C*, *SOS2*, *ACA2*, *ECA4*, and *MYB60,* in the wild type, are likely to be important regulatory changes that facilitate nocturnal stomatal opening and light period closure (Figure 4 and 8). As *MYB60* is required for stomatal opening in response to light in C_3_ Arabidopsis (Cominelli et al., 2005), it is particularly noteworthy that its transcript abundance in wild type *K. laxiflora* GC-enriched epidermal peels peaked at dusk, when stomata open in the wild type (Figure 8LL). Furthermore, in *rPPC1-B*, *MYB60* also had a dramatic 3- to 4-fold induction at dawn relative to the wild type, suggesting that high *MYB60* at the start of the light period may play a key role in the observed light period stomatal opening of the *rPPC1-B* line. It was also notable that *rPPC1-B* continued to have a peak of *MYB60* transcripts phased to dusk, as observed for the wild type, which correlated with the fact that *rPPC1-B* did open its stomata slightly throughout the dark period (Figure 4B). However, this nocturnal stomatal opening was futile in terms of atmospheric CO_2_ fixation, as *rPPC1-B* released respired CO_2_ from its leaves throughout the dark period (Figure 4A). Overall, *MYB60,* as well as several other mis-regulated guard cell signalling, ion channel and metabolite transporter genes (Figure 8 and Supplemental Figure 5S), represent key targets for future genetic manipulation experiments in transgenic *Kalanchoë* aimed at understanding the important regulators underpinning the inverse stomatal control associated with CAM.

### Informing biodesign strategies for engineering CAM into C_3_ crops

Efforts are underway to engineer CAM and its associated increased WUE into C_3_ species as a means to develop more climate-resilient crop varieties that can continue to fix CO_2_ and grow in the face of drought, whilst using water more wisely than C_3_ varieties (Borland et al., 2014; Borland et al., 2015; Lim et al., 2019). The data presented here for the *rPPC1* loss-of-function lines of *K. laxiflora* confirm experimentally the proposed core role of *PPC1* for efficient and optimised CAM. Our results also provide encouragement that the level of over-expression of a CAM-recruited *PPC1* gene introduced into an engineered C_3_ species may not need to be as high as the level found in extant obligate CAM species, because reducing *PPC1* levels to only 43 % of wild type activity in line *rPPC1-A* led to plants that were still capable of full CAM, and that fixed more CO_2_ than the wild type over 3 weeks of drought (Figure 4D).

The results presented here also emphasise that only certain sub-components of the core circadian clock are essential for the temporal optimisation of CAM in *Kalanchoë*, which in turn further simplifies the challenge of achieving correct temporal control of an engineered CAM pathway introduced into a C_3_ species. Transgenic manipulation of the expression and regulation of *CCA1-1*, *PRR3-7*, *PRR9* and *LNK3-like* in *Kalanchoë* using RNAi, over-expression and/ or CRISPR-Cas mediated gene editing, will allow the further refinement of the evolving model for the subset of core clock genes that form the transcription-translation feedback loop that underpins the temporal optimisation of CAM.

## MATERIALS AND METHODS

### Plant materials

*Kalanchoë laxiflora* were propagated clonally from leaf margin adventitious plantlets using the same clonal stock originally obtained from the Royal Botanic Gardens (RBG), Kew in 2008: accession number 1982-6028, which was originally kindly provided by the RBG-Kew Living Collection, but mis-identified under the name *K. fedtschenkoi*. Plants were grown initially in a heated transgenic glasshouse with supplementary lighting providing 16-h-light/ 8-h-dark and a minimum temperature of 20°C. Prior to all experiments, clonal populations of developmentally-synchronised plants were entrained for 7 days under 12-h-light/ 12-h-dark and/ or subsequently released into constant light and temperature conditions according to Boxall et al. (2017).

### Time course experiments

For light/dark time course experiments, opposite pairs of LP6 were collected every 4 h over a 12-h-light/12-h-dark cycle (12:12 LD), starting 2 h (02:00) after the lights came on at 00:00 h. Plants were entrained for 7-days in a Snijders Microclima MC-1000 growth cabinet set to 12-h-light (450 µmoles m^−2^ s^−1^), 25°C, 60 % humidity/ 12-h-dark, 15°C, 70 % humidity. For constant light, constant temperature, constant humidity (LL) free-running circadian time course experiments, plants were entrained under 12:12 LD conditions as above and switched to LL after a dark period. For LL, the constant conditions were: light 100 µmol m^−2^ s^−1^, temperature 15°C and humidity 70 %. LP6 were sampled every 4 h from three individual (clonal) plants, starting at 02:00 (2 h after lights on). All sampling involved immediate freezing of leaves in liquid nitrogen. Samples were stored at −80°C until use.

### Generation of transgenic *K. laxiflora* lines

An intron containing hairpin RNAi construct was designed to target the silencing of both copies of the CAM-associated *PPC1* gene in the tetraploid *K. laxiflora* genome (JGI Phytozome accession numbers: Kalax.0018s0056.1 and Kalax.0021s0061.1, which are the *K. laxiflora* orthologues of previously characterised *K. fedtschenkoi* CAM-associated *PPC1,* GenBank accession: KM078709). A 323 bp fragment was amplified from CAM leaf cDNA using high fidelity PCR with KOD Hot Start DNA Polymerase (Merck, Germany). The amplified fragment spanned the 3’ end of the *PPC1* coding sequence and extended into the 3’ untranslated region to ensure specificity of the silencing to both of the aforementioned CAM-associated *PPC1* gene copies. Alignment of the 323 bp region with the homologous regions from the six other plant-type *PPC* genes in the *K. laxiflora* genome demonstrated that none of the other *PPC* genes shared any 21 nucleotide stretches that were an exact match for the 323 bp *PPC1* RNAi fragment. Thus, the RNAi construct in the hairpin RNA binary vector used to generate the stable transgenic lines was predicted to silence only the two copies of the CAM-associated *PPC1,* and to be equally specific and efficient at silencing both copies. This specificity and equality of silencing efficiency was confirmed by the RT-qPCR data in Figure 1, as the qPCR primers targeted both copies of the CAM-associated *PPC1*, and so the quantitative signal is representative of the averaged signal for the transcript abundance of the two gene copies.

The primers used for the amplification of the *PPC1* gene fragment used to generate the hairpin RNAi binary construct were: *PPC1 RNAi F* 5’ CACCAAGCTACCAAGTGCCGGTG 3’, and *PPC1 RNAi R* 5’ CCCTCCTGCTGCTGCTGCTGC 3’. The PCR product was cloned into the pENTR/D Gateway-compatible entry vector, as described previously (Dever et al., 2015). Following confirmation of the correct sequence and orientation of the *PPC1* fragment in the ENTRY vector using Sanger sequencing, the pENTR/D *PPC1* RNAi clone was recombined into intron-containing hairpin RNAi binary vector pK7GWIWG2(II) (Karimi et al., 2002) using LR Clonase II enzyme mix (Life Technologies). *Agrobacterium*-mediated stable transformation of *K. laxiflora* was achieved using the method described previously for the very closely related species *K. fedtschenkoi* (Dever et al., 2015), with the following changes. For *K. laxiflora* transformation, the initial sterile leaf explants used for dipping in the *Agrobacterium tumefaciens* GV3101 carrying the engineered binary vector were generated by germinating surface-sterilised *K. laxiflora* seed on Murashige and Skoog medium with Gamborgs B-5 vitamins (Murashige and Skoog, 1962; Gamborg et al., 1968) and 3 % sucrose. 4- to 6-week-old seedlings raised in tissue culture were chopped into explants for transformation in a sterile laminar flow bench using a sterile scalpel. The explants were dipped in the *Agrobacterium* suspension carrying the *PPC1* RNAi binary vector, and co-cultivated and regenerated as described previously (Dever et al., 2015).

### High-throughput leaf acidity and starch content screens

Leaf acidity (as a proxy for leaf malate content) and leaf starch content were screened with leaf disc stains using chlorophenol red and iodine solution at both dawn and dusk as described by Cushman et al. (2008). For each transgenic line, leaf discs were sampled from LP6 in triplicate at 1 hour before dawn and 1 hour before dusk and stained in a 96 well plate format.

### Net CO_2_ exchange measurement

Gas exchange measurements were performed using a custom-built, 12-channel IRGA system (PP Systems, Hitchin, UK), which allowed individual environmental control (CO_2_/ H_2_O) and measurement of rates of CO_2_ uptake for each of 12 gas exchange cuvettes, with measurements collected every ∼18 min. The system was described in full by Dever et al. (2015), but was expanded here with the addition of a further 6 cuvettes, thereby doubling scope for replication and throughput. All experiments were repeated at least 3 times using 3 separate individual young plants (9-leaf-pairs), or detached LP6 from three separate clonal plants of each line. Representative gas exchange traces are shown. Replicated wild type and *rPPC1* lines were compared in neighbouring gas exchange cuvettes during each experimental run, such that the data are directly comparable between each line. As the entire gas exchange system was housed in a Snijders Microclima MC-1000 growth cabinet, all 12 gas exchange cuvettes were under identical conditions in terms of light intensity and temperature.

### Net CO_2_ exchange using the LI-COR 6400XT system

The gas exchange of mature CAM leaves (LP6) was measured over a 12-h-light, 25°C, 60 % humidity: 12-h-dark, 15°C, 70 % humidity cycle using an infra-red gas analyser (LI-6400XT, LI-COR, Inc) attached to a large CO_2_ gas cylinder. Data was logged every 10 minutes using an auto-program that tracked the light and temperature regimes of the growth cabinet.

### Leaf malate, starch and sucrose content

LP6 (full CAM in wild type) from mature plants were sampled into liquid nitrogen at the indicated times and stored at −80°C until use. The frozen leaf samples were prepared and assayed for malate and starch as described by Dever et al. (2015) using the published methods for assaying malate in an enzyme-linked spectrophotometric assay (Möllering, 1985), and starch (Smith and Zeeman, 2006). Sucrose, glucose, and fructose were assayed according to the manufacturer’s protocol (Megazyme Technologies).

### Chlorophyll Assays

Chlorophyll was assayed from mature greenhouse grown plants from leaf pairs 2-7. Chlorophyll was extracted twice from 0.5 cm diameter leaf discs in 2 ml 80 % (v/v) acetone and homogenised in a bead beater (PowerLyzer 24; Mo-Bio, Inc). The tubes were centrifuged at full speed in a bench top microfuge at 4°C for 2 min, and the supernatants were combined and transferred to a new tube and protected from the light. Absorbance was read at 663 nm and 645 nm, and chlorophyll contents were calculated according to the published method (Arnon, 1949).

### Total RNA isolation and RT-qPCR

Total RNA was isolated from 100 mg of frozen, ground leaf tissue using the Qiagen RNeasy kit (Qiagen, Germany) following the manufacturer’s protocol with the addition of 13.5 µL 50 mg mL^−1^ PEG 20000 to the 450 µL RLC buffer used for each extraction. cDNA was synthesised from the total RNA using the Qiagen Quantitect RT kit according to the manufacturer’s instructions (Qiagen, Germany). The resulting cDNA was diluted 1:4 with molecular biology grade water prior to use in RT-qPCR. Transcript levels were determined using the SensiFAST SYBR No Rox kit (Bioline) in an Agilent MX3005P qPCR System Cycler. The results for each target gene transcript of interest were normalized to the reference gene *THIOESTERASE/THIOL ESTER DEHYDRASE-ISOMERASE SUPERFAMILY PROTEIN* (*TED*; Kalax.0134s0055.1 and Kalax.1110s0007.1; Arabidopsis orthologue AT2G30720.1). Gene expression in a pool of RNA generated from LP6 samples collected every 4 hours over a 12-h-light/ 12-h-dark cycle was set to 1. Primers for RT-qPCR analyses are listed in supplemental table S1.

### Immunoblotting

Total protein extracts of *K. laxiflora* leaves were prepared according to Dever et al. (2015). One-dimensional SDS-PAGE and immunoblotting of leaf proteins was carried out following standard methods. Blots were developed using the ECL system (GE Healthcare, UK). Immunoblot analysis was carried out using antisera to PPC raised against purified CAM leaf PPC from *K. fedtschenkoi,* kindly supplied by Prof. H.G. Nimmo, University of Glasgow (Nimmo et al., 1986), and the phosphorylated form of PPC raised against a phospho-PPC peptide from barley, and kindly supplied by Prof. Cristina Echevarría, Universidad de Sevilla, Spain (González et al., 2002; Feria et al., 2008). The PPDK antibody was raised against maize (*Zea mays*) C_4_ PPDK, and the phospho-PPDK antibody was raised against a synthetic phospho-peptide spanning the PPDK phosphorylation site in the *Z. mays* C_4_ PPDK. Both PPDK antibodies were kindly provided by Prof. Chris J. Chastain, University of Minnesota, Moorhead, USA (Chastain et al., 2000; Chastain et al., 2002).

### Growth measurements

Mature plants of wild type and the two *rPPC1* lines were grown from developmentally synchronized clonal leaf plantlets in greenhouse conditions for 4 months. At the start of the drought treatment all the plants were watered to full capacity. Water was withheld from 10 replicate plants of each line for 28 days, and 10 plants were maintained well-watered over the same period. The plants were harvested as separated above-ground (shoot) and below-ground (root) tissues, weighed to determine fresh weight, and then dried in an oven at 60°C until they reached a constant dry weight.

### PPC assays

Frozen leaf tissue was ground in liquid nitrogen with a small quantity of acid washed sand and the relevant enzyme specific extraction buffer (approximately 1 g tissue to 3 ml of extraction buffer). Extracts were prepared and PPC assays were performed according to the extraction, desalting and assay buffer conditions described previously (Dever et al., 2015).

### Determination of the apparent K_i_ of PPC for L-malate in rapidly desalted leaf extracts

The apparent K_i_ of PPC for L-malate was determined using leaf extracts that were rapidly desalted as described by Carter et al. (1991). LP6 were collected at 10:00 (2 h before the end of the 12-h-light period), and 18:00 (middle of the 12-h-dark period) from three biological replicates of wild type, *rPPC1-A* and *rPPC1-B*. The apparent K_i_ of PPC activity for feedback inhibition by L-malate was determined according to the method described by Nimmo et al. (1984), with the modifications to the range of L-malate concentrations added to the assays as described by Boxall et al. (2017).

### PPDK assays

PPDK assays were performed using the extraction and assay buffers described previously (Kondo et al., 2000; Dever et al., 2015), with the addition of NADH and G6P (Salahas et al., 1990), and Cibercron Blue (Burnell and Hatch, 1986). Briefly, 0.3 g of powdered leaf tissue that had previously been ground to a fine powder in liquid nitrogen was extracted in 1 ml of ice-cold extraction buffer containing 100 mM Tris pH 8.0, 10 mM DTT, 1 mM EDTA, 1 % Triton, 2.5 % w/v PVPP, 2 % PEG-20000, 10 mM MgCl_2_, 1 mM PMSF, 2 µM orthovanadate, and 10 µM Cibercron Blue, by grinding with a small quantity of acid washed sand in a pestle and mortar. Extracts were vortexed for 30 s and the pH was adjusted to pH 8.0. Extracts were then placed on ice for 10 mins before spinning them at full speed in a benchtop microfuge at 4°C. The supernatant (500 µl) was desalted using PD minitrap Sephadex-G25 columns (GE Healthcare). The desalting buffer contained: 100 mM Tris-HCl pH 8.0, 10 mM MgCl_2_, 10 mM DTT, 0.1 mM EDTA, 10 µM Cibercron Blue. The desalted extracts were assayed in a plate reader at 340 nm using the following assay buffer: 100 mM Tris pH 8.0, 5 mM DTT, 10 mM MgCl_2_, 1.25 mM Pyruvic acid, 2.5 mM NaHCO_3_, 2.5 mM K_2_HPO_4_, 0.25 mM NADH, 2 U Malate dehydrogenase, 6 mM glucose 6-phosphate. The reactions were started by the addition of 0.2 U of phosphoenolpyruvate carboxylase followed by 1.25 mM ATP (pH8.0).

### Delayed fluorescence (DF) measurements

The imaging system for DF was identical to the luciferase and delayed fluorescence imaging system described previously (Gould et al., 2009) with the exception of the CCD camera (Retiga LUMO™ CCD Camera; Qimaging). DF was quantified using image analysis software Imaris (Bitplane) to measure mean intensity for specific regions within each image. Background intensities were calculated for each image and subtracted to calculate a final DF value for each image (Gould et al., 2009).

### DF rhythm analysis

*K. laxiflora* plants were grown in greenhouse conditions for 4 months and then entrained in 12-h light (450 µmoles m^−2^ s^−1^), 25°C, 60 % humidity/ 12-h dark, 15°C, 70 % humidity cycles in a Snijders Microclima MC-1000 (Snijders Scientific) growth cabinet for 7 d as described previously (Dever et al., 2015). At dawn on the 8th day, 1.5-cm leaf discs were punched from each of LP6 for three biological replicates (i.e., 7 leaf discs from each biological replicate, totalling 21 leaf discs per line) and placed on 0.3 % Phytoagar (Duchefa Biochemie) on a 10-cm square Petri dish. The Petri dish was left under 12-h light/25°C: 12-h dark/15°C for a further 24 h to synchronise the leaf discs. At subjective dawn the Petri dish was placed in the imaging system at 14°C in constant red-blue light (LL). DF images were collected every hour for 120 h as described previously (Gould et al., 2009; Boxall et al., 2017). The DF images were processed as described by Gould et al. (2009). The luminescence was normalized by subtracting the Y value of the best straight line from the raw Y value. Biodare was used to carry out fast Fourier transform (nonlinear least square) analysis and spectral resampling on each DF time-course series using the time window from 24 to 120 h in order to generate period estimates and calculate the associated relative amplitude error (RAE) (Moore et al., 2014; Zielinski et al., 2014).

### Accession numbers

Sequence data associated with this article are available via the JGI Phytozome portal for the *K. laxiflora* genome: https://phytozome.jgi.doe.gov ***Kalanchoë laxiflora* v1.1.** and the specific gene IDs for each gene measured in this work are provided in Supplemental Table 1.

## ACKNOWLEDGEMENTS

We thank Prof. Hugh Nimmo (University of Glasgow, UK), Prof. Cristina Echevarria (Universidad de Sevilla, Spain) and Prof. Chris Chastain (Minnesota State University-Moorhead, USA) for providing the antibodies used in this study. This work was supported by the U.S. Department of Energy (DOE) Office of Science, Genomic Science Program under Award Number DE-SC0008834, and in part by the Biotechnology and Biological Sciences Research Council, U.K. (BBSRC grant no. BB/F009313/1 awarded to J.H.). The contents of this article are solely the responsibility of the authors and do not necessarily represent the official views of the DOE.

## AUTHOR CONTRIBUTIONS

J.H. and S.F.B. designed the research. S.F.B. performed all of the experiments except some of the tissue culture work for the regeneration of the transgenic lines was performed by N.K., and the statistical analysis of the delayed fluorescence data was performed by P.J.D.G. J.K. generated the RNAi binary construct for *PPC1*. L.V.D. performed the PPDK assays. J.L.W. grew and maintained the plants and helped with grinding and processing of leaf samples. P.J.D.G. helped S.F.B. with the delayed fluorescence experiments and the analysis and interpretation of the associated data. S.F.B. and J.H. wrote the manuscript.

---

**Supplemental Table S1:**
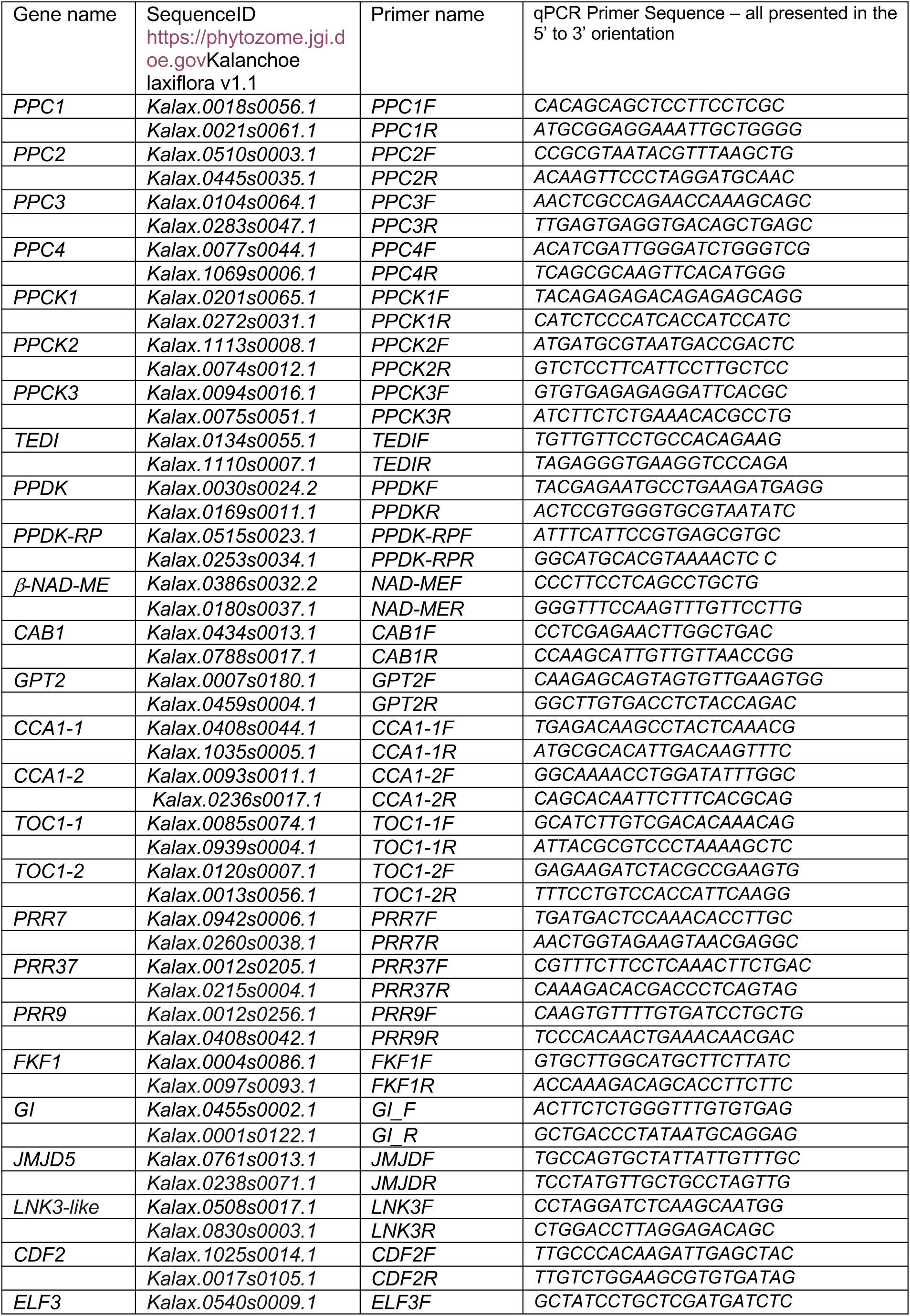
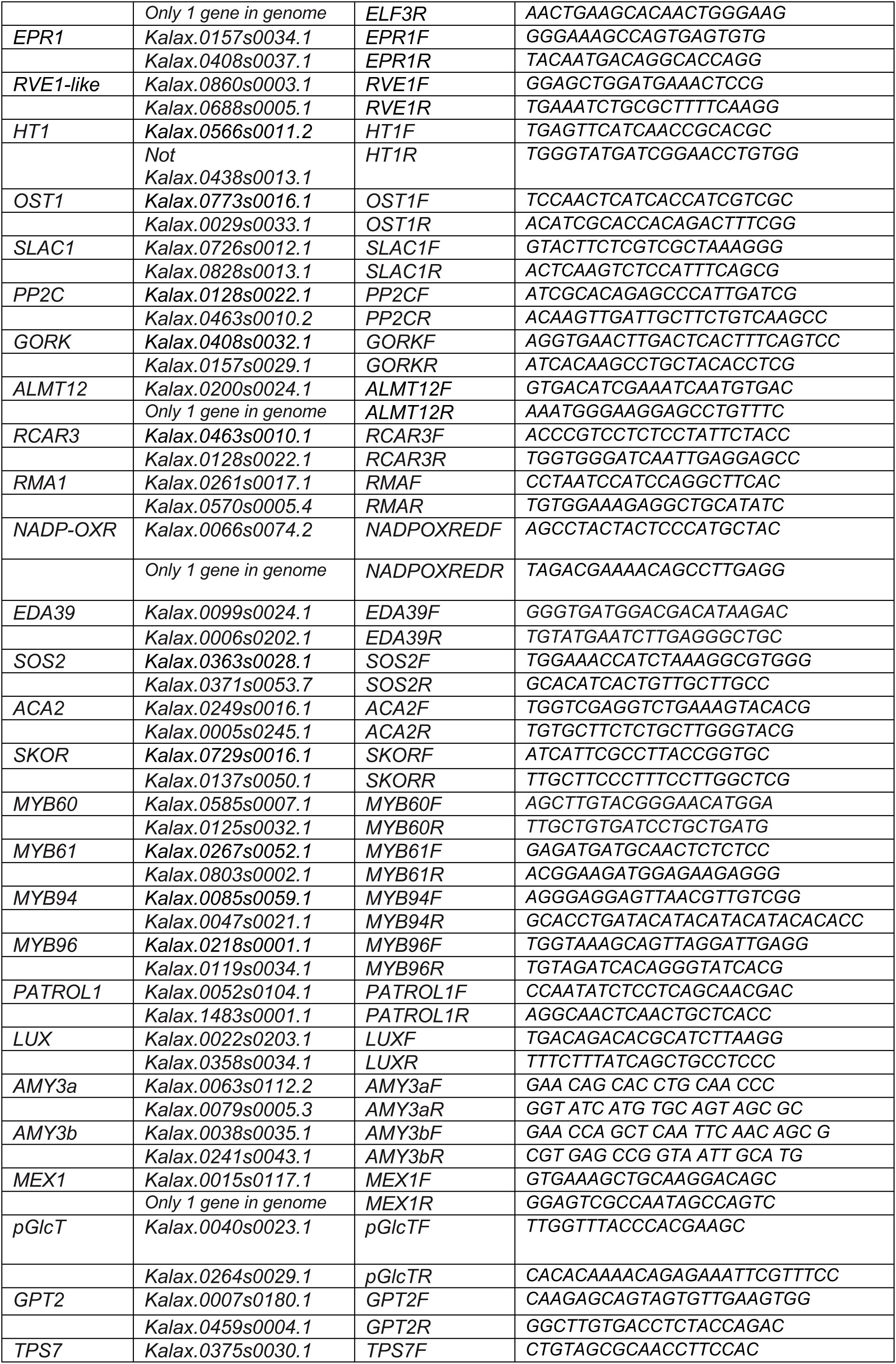
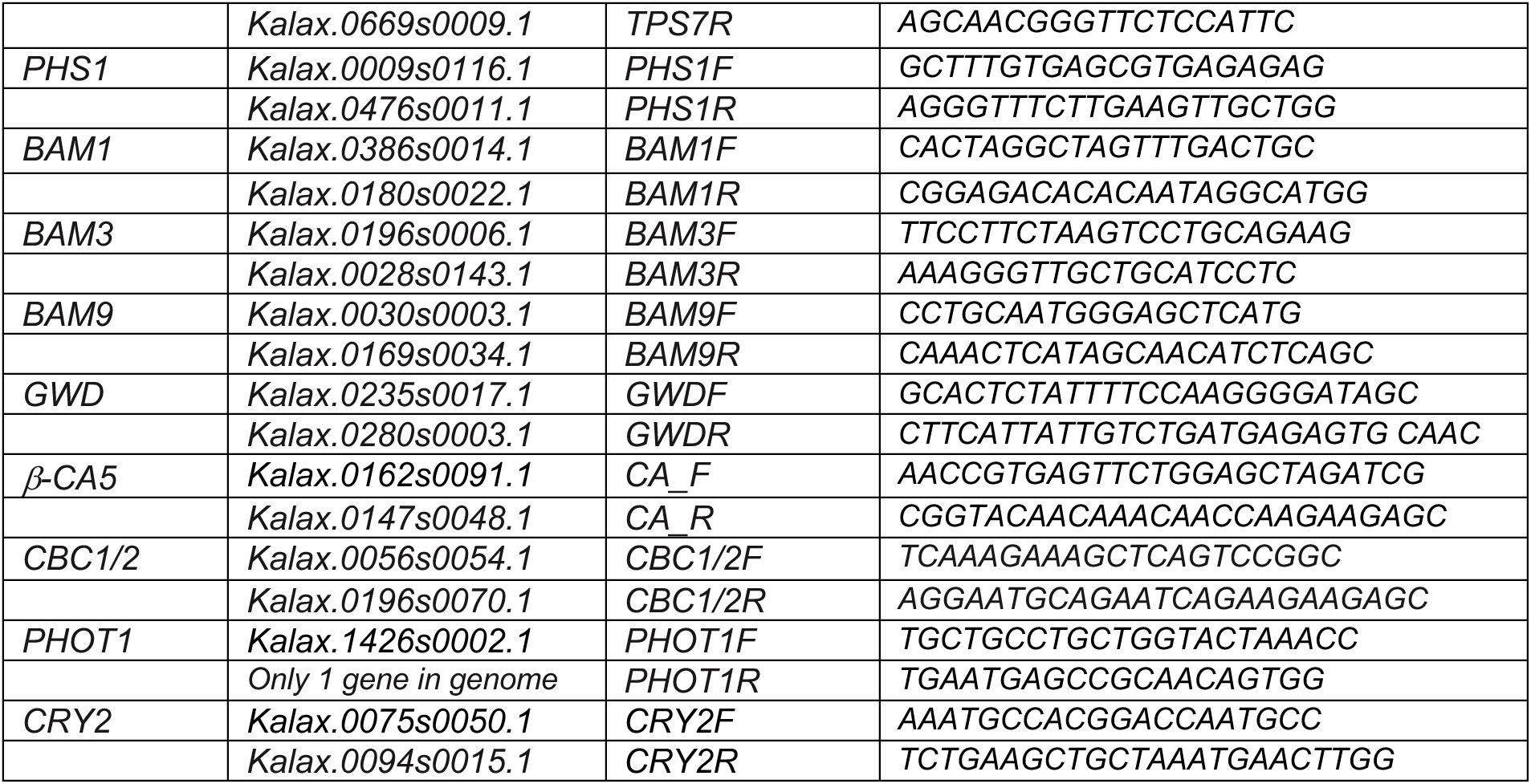
Primers used for RT-qPCR

**Supplemental Table 2:** Calculations for the cumulative total CO_2_ fixation by each line during the 27-day drought-stress and re-watering experiment presented in Figure 4D.

|  | Wild Type<br>d1-d27 | *rPPC1-A*<br>d1-d27 | *rPPC1-B*<br>d1-d27 | Wild type<br>drought<br>period<br>d1-d22 | *rPPC1-A*<br>drought<br>period<br>d1-d22 | *rPPC1-B*<br>drought<br>period<br>d1-d22 |
| --- | --- | --- | --- | --- | --- | --- |
| Area of Positive Peaks | 25630 | 26262 | 24360 | 19911 | 20560 | 17516 |
| Std. Error | 172.1 | 138.8 | 148 | 155.1 | 128.6 | 142.1 |
| 95% Confidence Interval | 25292 to 25967 | 25990 to 26534 | 24070 to 24650 | 19607 to 20214 | 20307 to 20812 | 17238 to 17795 |
| Area of Negative Peaks | 1514 | 1588 | 5529 | 1343 | 1489 | 3533 |
| Std. Error | 42.73 | 36.83 | 71.61 | 27.05 | 24.19 | 26.68 |
| 95% Confidence Interval | 1430 to 1598 | 1516 to 1661 | 5389 to 5670 | 1290 to 1396 | 1442 to 1536 | 3481 to 3586 |
| Total Area | 27143 | 27851 | 29889 | 21253 | 22048 | 21050 |
| Std. Error | 177.3 | 143.6 | 164.4 | 157.4 | 130.9 | 144.6 |
| 95% Confidence Interval | 26796 to 27491 | 27569 to 28132 | 29567 to 30212 | 20945 to 21562 | 21792 to 22305 | 20766 to 21333 |
| Net Area | 24116 | 24674 | 18831 | 18568 | 19071 | 13983 |
| Std. Error | 177.3 | 143.6 | 164.4 | 157.4 | 130.9 | 144.6 |
| 95% Confidence Interval | 23768 to 24463 | 24392 to 24955 | 18509 to 19153 | 18259 to 18876 | 18814 to 19327 | 13700 to 14267 |
| Total Peak Area | 27143 | 27851 | 29889 | 21253 | 22048 | 21050 |
| Std. Error | 177.3 | 143.6 | 164.4 | 157.4 | 130.9 | 144.6 |
| 95% Confidence Interval | 26796 to 27491 | 27569 to 28132 | 29567 to 30212 | 20945 to 21562 | 21792 to 22305 | 20766 to 21333 |

## Supplemental Results Text

### Diel Regulation of GC-Metabolism Genes in GC-Enriched Epidermal Peels of rPPC1-B

In *rPPC1-B,* stomatal opening shifted to the light period (Figure 4A to 4C). It was thus important to investigate whether the temporal pattern of GC-specific regulatory genes was rescheduled in *rPPC1-B*. RNA was isolated from epidermal peels separated from mature leaves (LP6 to LP8). Peels were enriched with intact stomatal GCs engaged in the CAM pattern of opening and closing in the wild type.

*PPC1* and *PPC2* transcripts were low in epidermal peels (Supplemental Figure S5A and S5B) when compared to whole leaves (Figure 1A). *PPC3* and *PPC4* were more abundant transcripts in wild type epidermal peels (Figure S5C and S5D) relative to whole leaves (Figure 1C and 1D). Furthermore, *PPC3* and *PPC4* were up-regulated in epidermal peels of *rPPC1-B* compared to the wild type, and their transcript abundance peaked 4 h into the dark, whereas in the wild type both transcripts peaked at the end of the dark period or dawn (Figures S5A to S5D).

CAM-specific *PPCK1* was up-regulated in *rPPC1-B* epidermal peels compared to wild type, whereas *PPCK2*, which does not currently have a proposed role in the regulation of CAM *PPC1* in the mesophyll, was unchanged (Figure S5E and S5F). Both *PPCK2* and *PPCK3* were more abundant than *PPCK1* in epidermal peels, suggesting their encoded proteins may function to regulate the activity of the GC PPC(s) (Figure S5F and S5G). Most strikingly, *PPCK3* was up-regulated ∼5-fold at 4 h after lights-on in *rPPC1-B* epidermal peels relative to the wild type (Figure S5G).

In C_3_ plants, starch is degraded before dawn to fuel stomatal opening (Blatt, 2016). In Arabidopsis GCs, *BAM1* encodes the major starch degrading enzyme (Valerio et al., 2010; Prasch et al., 2015; Horrer et al., 2016; Santelia and Lunn, 2017) that functions with AMY3 to mobilize starch at dawn, releasing maltose in the chloroplasts (Horrer et al., 2016). In *rPPC1-B* epidermal peels*, GWD* was halved relative to wild type at its peak, 8 h into the 12 h light period (Figure S5H). *AMY3a* and *AMY3b* were up-regulated at dusk and in the second half of the light period, respectively (Figure S5I and S5J). *BAM1* was up-regulated at dawn compared to wild type, and peaked with a very similar transcript abundance to wild type at 20:00 h, 8 h into the dark period (Figure S5K), whereas *BAM3* was ∼3-fold lower in *rPPC1-B* relative to the wild type at the time of its nocturnal peak, which also occurred 8 h into the dark (Figure S5L). *BAM9* in *rPPC1-B* rose later than the wild type, lagging behind the wild type at dusk and 4 h into the dark, and it stayed ∼5-fold higher at dawn and 4 h into the light, when the wild type reached its daily trough (Figure S5M). *BAM9* in Arabidopsis is predicted to be catalytically inactive and has no known function to date (Monroe and Storm, 2018). The peak of *PHS1* was delayed by 4h peaking at the light/ dark transition (Figure S5N), and *MEX1* was down-regulated in epidermal peels of *rPPC1-B,* peaking at 08:00 in the light, 4 h after the wild type peak (Figure S5O).

### Diel Regulation of Core Circadian Clock genes in Epidermal Peels of *rPPC1-B*

The circadian clock genes *PRR3/7* and *PRR7* were up-regulated in epidermal peels of *rPPC1-B* relative to wild type (Figure S5P and S5Q), with a similar diel pattern to that measured in whole CAM leaves (Figure 5S and 5T). *PRR3/7* peaked 4 h earlier in *rPPC1-B*, whereas *PRR7* transcript levels were greater throughout the light period in *rPPC1-B* (Figure S5P and S5Q). *LUX* is a component of the morning transcriptional feedback circuit within the clock, and acts as a transcription factor that directly regulates the expression of *PRR9* by binding specific sites in its promoter (Helfer et al., 2011). In epidermal peels, *LUX was* down regulated >7-fold in *rPPC1-B* compared to the wild type at its peak, 4 h into the light, and also peaked 4 h later than the wild type (Figure S5R).

### Diel Regulation of GC-Signalling Genes in GC-Enriched Epidermal Peels of rPPC1-B

GCs perceive CO_2_ and regulate stomatal aperture via ABA signalling and reactive oxygen species, with low CO_2_ mediating stomatal opening and high CO_2_ causing closure (Chater et al., 2015). NADP oxidoreductase encodes an NADP-binding Rossman-fold super-family protein that was only detected in epidermal peels in *Kalanchoë* (Boxall, Dever, Kadu and Hartwell, unpublished observation). The enzyme encoded by this gene may function to generate the H_2_O_2_ burst induced by ABA as part of the stomatal closure signalling pathway (Daszkowska-Golec and Szarejko, 2013). This transcript was up-regulated ∼3-fold in *rPPC1-B* at dawn (Figure S5S).

In Arabidopsis, transcription factors *MYB94* and *MYB96* function together in the activation of cuticular wax biosynthesis under drought stress (Lee et al., 2016). *MYB96* may also be involved in the response to drought stress in Arabidopsis through ABA signalling that mediates stomatal closure via the *RD22* pathway (Seo et al., 2011). Both *MYB94* and *MYB96* were induced by as much as 3-fold in *rPPC1-B* compared to wild type (Figure S5T and S5U). Specifically, *MYB94* transcript levels rose 8 h earlier than they did in the wild type and were already close to peak levels by the light-to-dark transition, whereas wild type *MYB94* levels peaked sharply 8 h into the dark period (Figure S5T). *MYB96* transcript levels were induced relative to the wild type at both dawn and dusk, but the peak of *MYB96* at dawn represented the largest fold-change relative to the wild type (Figure S5U).

**Figure S1.**
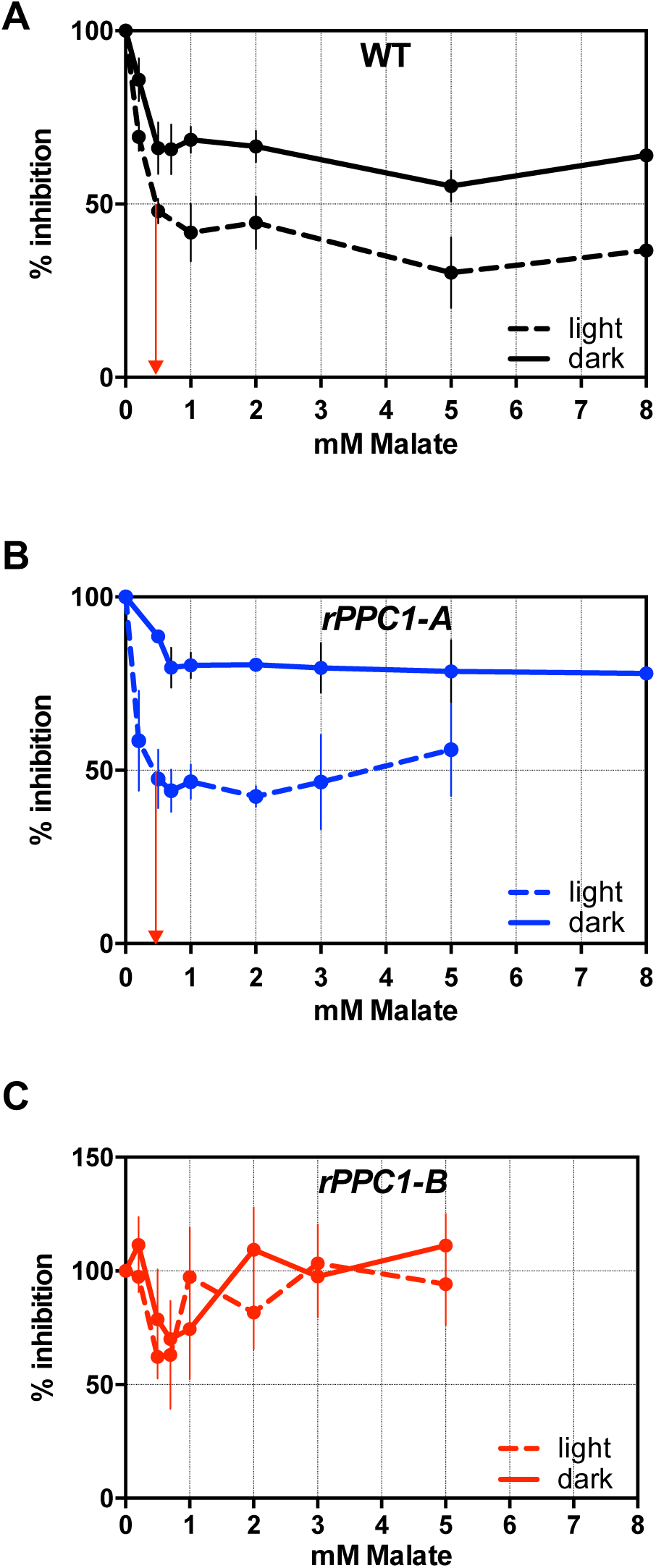
Malate sensitivity assays for PPC activity measured in rapidly desalted extracts prepared from leaf pair 6. **(A)**, wild type, **(B)**, *rPPC1-A* and **(C)**, *rPPC1-B.* Desalted extracts were prepared from CAM leaves (leaf pair 6; LP6) using plants entrained in 12-h-light/ 12-h-dark. Inhibition of PPC by malate is shown in LP6 taken at 6 h into light and 10h into the dark. Black data are for the wild type, blue data *rPPC1-A,* and red data *rPPC1-B*.

**Figure S2.**
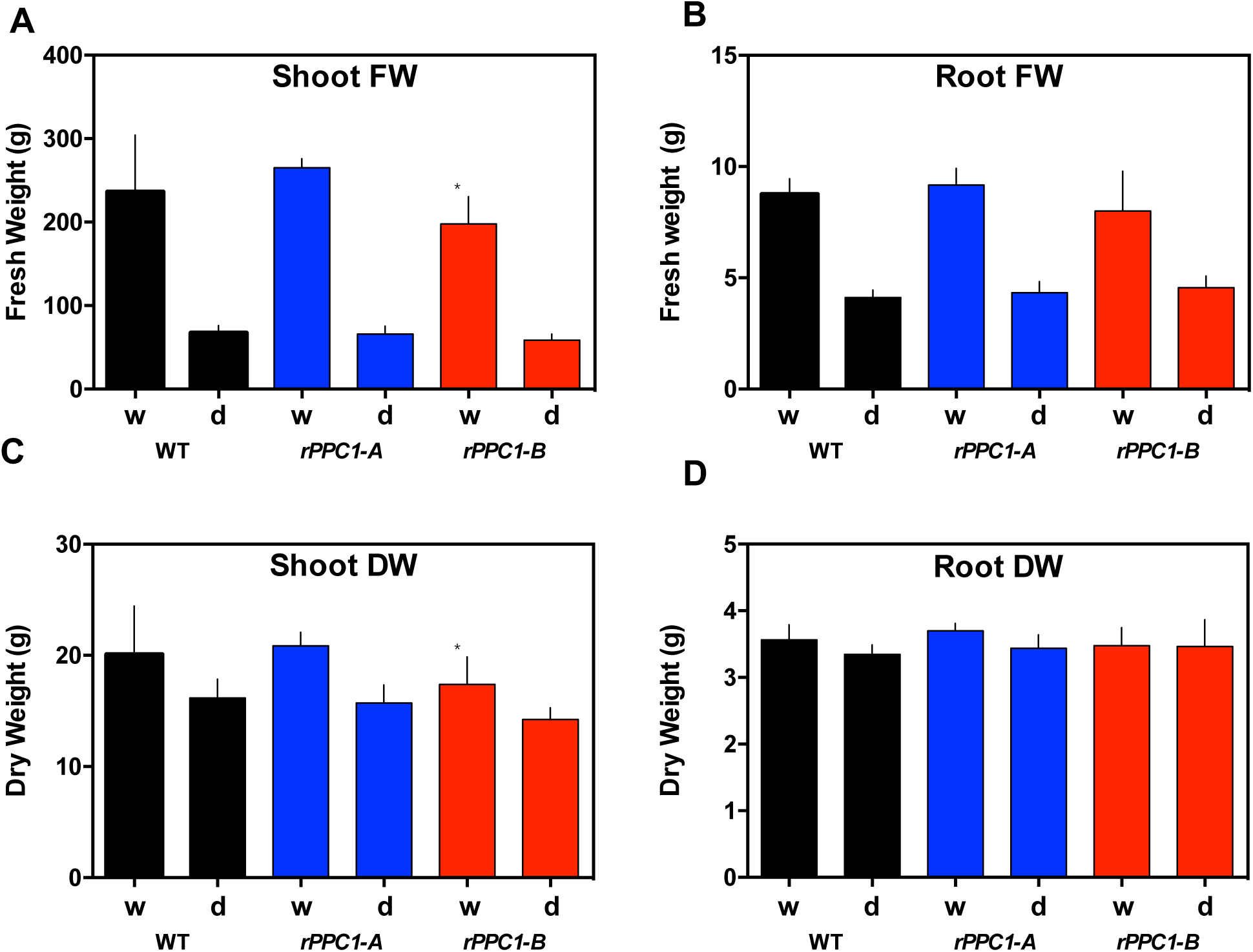
Impact of Loss of PPC Activity on Vegetative Yield during Growth under Well-Watered (w) and Drought-Stressed (d) Conditions. **(A)**, shoot fresh weight, **(B)**, root fresh weight, **(C)**, shoot dry weight, and **(D)**, root dry weight. Fresh and dry weight of above ground biomass (shoot) and below-ground tissues (roots) at maturity following 138 d of growth under glassshouse conditions. Wild type (WT), *rPPC1-A and rPPC1-B* were measured under both well-watered and drought-stressed conditions. n = 6 plants; error bars represent standard errors. Black data are for the wild type, blue data *rPPC1-A,* and red data *rPPC1-B*. Asterisks indicate significant difference from the wild type based on Student’s t-test: *Shoot FW p = 0.026; Shoot DW p= 0.026, n = 10 developmentally synchronised clonal plants of each line.

**Figure S3.**
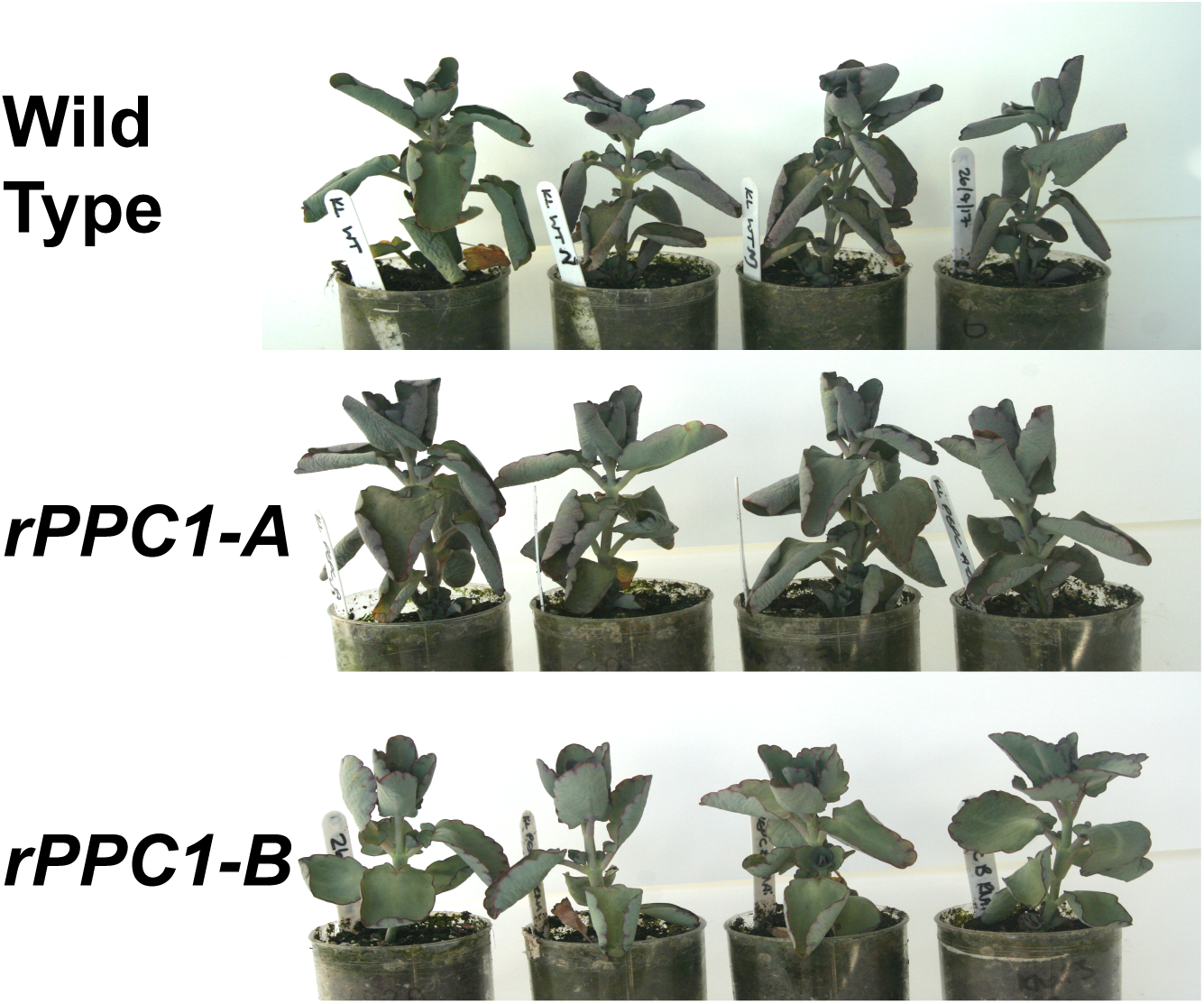
Photographs Demonstrating the Visual Appearance of Small Plants after 23 d Drought Prior to Re-watering. Wild type plants with normal levels of *PPC1* had visibly curled leaves after 23 d drought, whereas *rPPC1-B* leaves look less curled. Plants were removed from the gas exchange cuvettes after 23 d of drought, prior to re-watering. Plants were returned to the gas exchange cuvettes after re-watering for additional CO_2_ uptake measurements to be collected. The gas exchange data for these plants is presented in Figure 4D.

**Figure S4.**
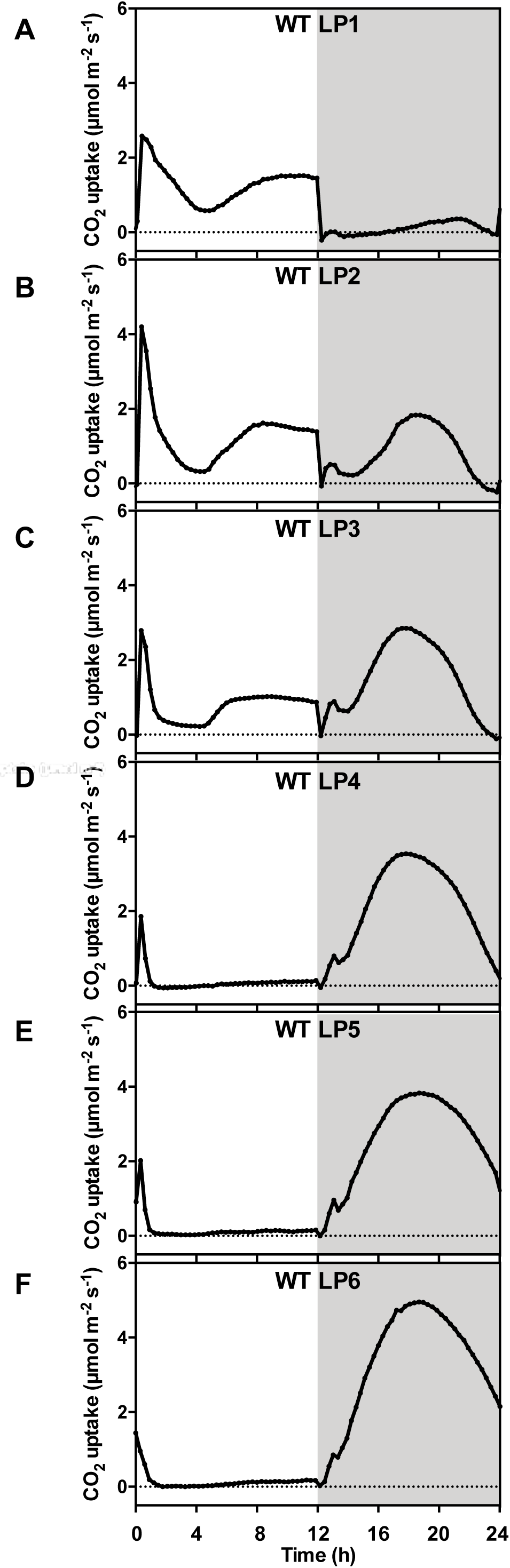
Impact of Leaf Age on the Development of the Characteristic 24 h Light/ Dark Pattern of CAM-Associated CO_2_ Exchange. Gas exchange profiles over a 24 h light/ dark cycle (12-h-light/ 12-h-dark; 25 °C: 15 °C; 60 %/ 70 % humidity) using detached leaves of increasing ages with their petioles placed in distilled water. (A), leaf pair 1 (youngest); (B), leaf pair 2; **(C)**, leaf pair 3; **(D)**, leaf pair 4; **(E)**, leaf pair 5, and **(F)**, leaf pair 6 (oldest).

**Figure S5.**
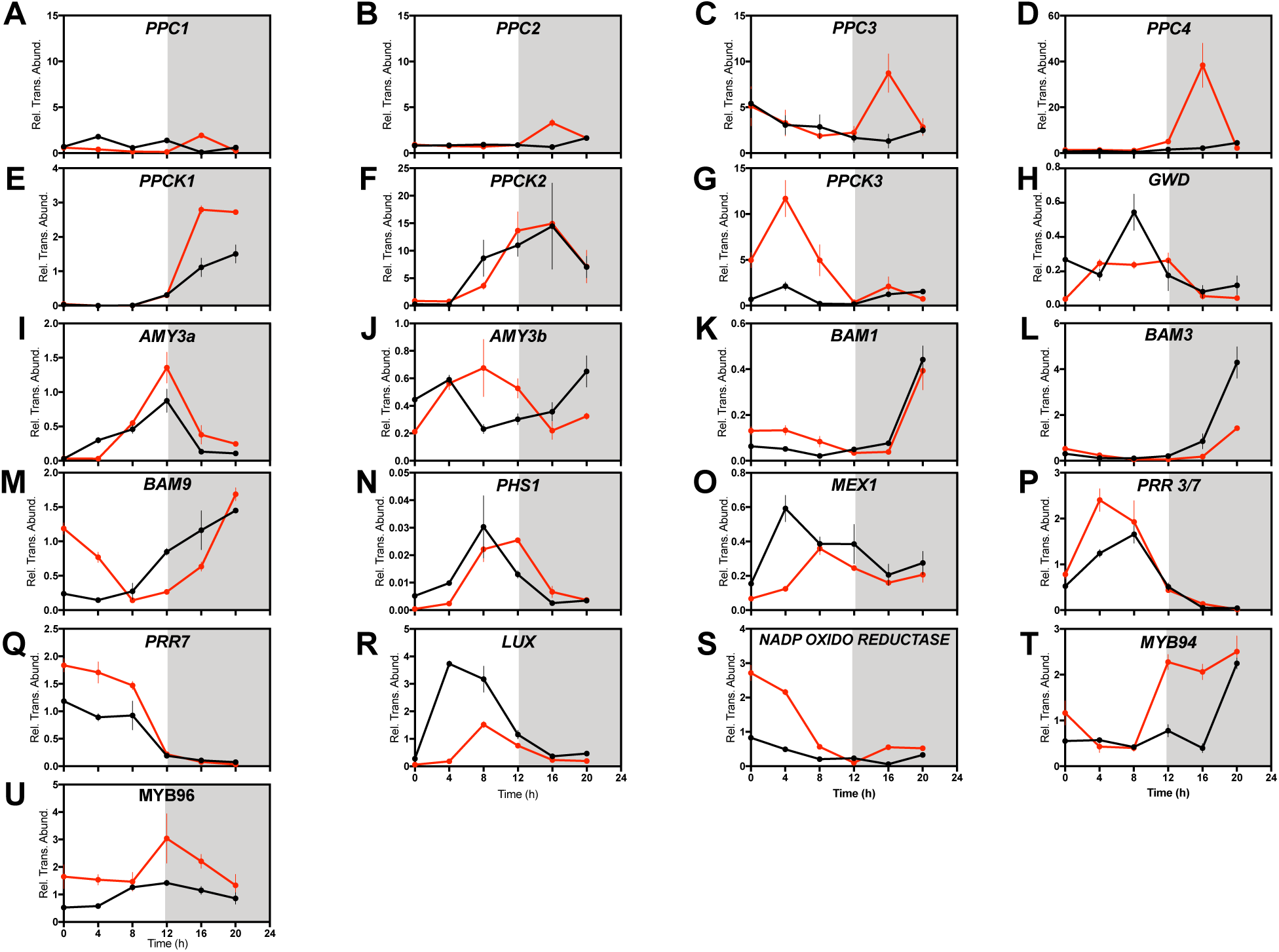
Impact of the Loss of PPC activity on the Light/ Dark Regulation of the Transcript Abundance of Stomatal Genes in the Epidermis. Plants were entrained under 12-h-light/ 12-h-dark for 7 days prior to sampling. Epidermal peel samples were separated from leaf pairs 6, 7 and 8, with samples collected every 4h starting at 02:00, 2 h into the 12-h-light period. Each biological sample represents a pool of epidermal peels taken from 6 leaf pair 6 leaves from 3 clonal stems of each line. Each peel was frozen in liquid nitrogen immediately after it was taken and pooled later. RNA was isolated and was used in RT-qPCR. A thioesterase/thiol ester dehydrase-isomerase superfamily gene (*TEDI*) was amplified as a reference gene from the same cDNAs. Gene transcript abundance data represent the mean of 3 technical replicates for biological triplicates, and was normalized to the reference gene *(TEDI)*; error bars represent the standard error. In all cases, plants were entrained under 12-h-light/ 12-h-dark cycles prior to release into LL free-running conditions. Black data are for the wild type and red data *rPPC1-B.* (A), *PPC1*; **(B)**, *PPC2*; **(C)**, *PPC3*; **(D)**, *PPC4*; **(E)**, *PPCK1*; **(F)**, *PPCK2*; **(G)**, *PPCK3*; **(H)**, *GWD*; **(I)**, *AMY3a*; **(J)**, *AMY3b*; **(K)**, *BAM1*; **(L)**, *BAM3*; **(M)**, *BAM9*; **(N)**, *PHS1*; **(O)**, *MEX1*; **(P)**, *PRR37*; **(Q)**, *PRR7*; **(R)**, *LUX*; **(S)**, *NADP OXIDOREDUCTASE*; **(T)**, *MYB94*; and (U), *MYB96*.

